# TRAPID 2.0: a web application for taxonomic and functional analysis of *de novo* transcriptomes

**DOI:** 10.1101/2020.10.19.345835

**Authors:** François Bucchini, Andrea Del Cortona, Łukasz Kreft, Alexander Botzki, Michiel Van Bel, Klaas Vandepoele

## Abstract

Advances in high-throughput sequencing have resulted in a massive increase of RNA-Seq transcriptome data. However, the promise of rapid gene expression profiling in a specific tissue, condition, unicellular organism, or microbial community comes with new computational challenges. Owing to the limited availability of well-resolved reference genomes, *de novo* assembled (meta)transcriptomes have emerged as popular tools for investigating the gene repertoire of previously uncharacterized organisms. Yet, despite their potential, these datasets often contain fragmented or contaminant sequences, and their analysis remains difficult. To alleviate some of these challenges, we developed TRAPID 2.0, a web application for the fast and efficient processing of assembled transcriptome data. The initial processing phase performs a global characterization of the input data, providing each transcript with several layers of annotation, comprising structural, functional, and taxonomic information. The exploratory phase enables downstream analyses from the web application. Available analyses include the assessment of gene space completeness, the functional analysis and comparison of transcript subsets, and the study of transcripts in an evolutionary context. A comparison with similar tools highlights TRAPID’s unique features. Finally, analyses performed within TRAPID 2.0 are complemented by interactive data visualizations, facilitating the extraction of new biological insights, as demonstrated with diatom community metatranscriptomes.

## INTRODUCTION

The advent of low-cost RNA sequencing (RNA-Seq) has led to a steep increase in the number of available transcriptomes for which no corresponding reference genome sequence is yet available. Sequencing and assembly often remain more cost-effective for *de novo* transcriptomes than for whole genomes, especially for organisms featuring complex and large genomes or for species that cannot be cultured in laboratory conditions (1). As the cost and labor to go from a low-quality fragmented genome assembly to a high-quality chromosome level assembly can be prohibitively high, a high-quality transcriptome can represent a valuable alternative to a low-quality genome. Although a transcriptome does not provide the same opportunities for downstream analyses as a fully sequenced and assembled genome, it still suits multiple purposes. Examples include studying the presence or absence of specific gene functions or pathways, as well as unraveling taxonomic relationships between sets of distantly related species (2–6).

Protists are unicellular eukaryotes that encompass the majority of the eukaryotic biodiversity (7,8), but for which genome-based studies are lagging behind. These organisms show a wide range of nutritional modes and have a major impact on the ecology and chemistry of their ecosystems. The size and complexity of eukaryotic genomes makes genome sequencing a daunting task, resulting in the widespread adoption of transcriptomic approaches to characterize the gene content of unicellular eukaryotes (9,10). Apart from transcriptome reconstruction, each transcript can also be quantified according to whether it is expressed in certain environmental conditions. As such, transcriptomic studies of cultures and natural assemblages of protists are revealing complex metabolic responses to environmental conditions (11). As a metatranscriptome collects all RNA transcripts present in a sample composed of a community of microorganisms, it represents a molecular readout of the global transcriptional gene activity present in a specific environment or in response to environmental perturbations such as nutrient availability, the presence of other (non-) parasitic species, or abiotic effects. Nevertheless, sufficient sequencing depth as well as the presence of high-quality and complete reference data sets are imperative for successful gene annotation of complex metatranscriptomes (12). Metatranscriptome sequence analysis can not only shed light on a species’ success in specific natural communities, but disentangling taxonomic information and gene functions encoded by specific transcripts allows for the study of the functional or metabolic contribution of individual species to complex environments (13–17).

Here we present TRAPID 2.0, the next iteration of the TRAPID (18) platform for fast and accurate transcriptome analysis, utilizing comparative genomics approaches with a series of high-quality reference genome databases. This new version features numerous improvements on both the backend and frontend: a reworked processing pipeline, updated reference databases, and a redesigned web application, together with a wide variety of new analytical features. Due to the improved processing speed and the integration of protein-based taxonomic sequence classification, TRAPID 2.0 now also supports the analysis of assembled (meta)transcriptomes showing different prokaryote and/or eukaryotic compositions.

## MATERIAL AND METHODS

### Collection of transcriptomes from unicellular eukaryotes

Transcriptomes of 34 samples from the Marine Microbial Eukaryotic Transcriptome Sequencing Project (MMETSP) re-assembled data set (9,19) were retrieved from http://doi.org/10.5281/zenodo.1212585 (version 2; see Supplementary Table S1 for a complete list). Selected ciliate MMETSP samples correspond to organisms reported to use the ciliate nuclear genetic code in the NCBI taxonomy.

### Taxonomic classification of transcripts

Taxonomic classification of transcript sequences is performed using Kaiju (20), run in MEM mode, using the parameters ‘-x -m 11’ (filtering of low-complexity sequences, minimum match length of 11 amino acids). The reference index used during the taxonomic classification consists of all sequences from eukaryotes, bacteria, archaea, and viruses in the NCBI non-redundant protein database (downloaded 5^th^ of September 2019). Because Kaiju’s index needs to be loaded in memory, we run Kaiju using a split index to keep memory usage reasonable and ensure we can face the perpetual data growth of the NCBI non-redundant protein database without major hurdles. The split results of Kaiju are merged using the same selection criteria as those used by Kaiju’s algorithm (i.e. keeping the longest match from all the splits, or the lowest common ancestor in case equally long matches were found). Benchmark experiments confirmed that splitting the index results in nearly identical performance compared to a regular Kaiju MEM run (Supplementary Note S1; Supplementary Figure S1). The ‘Krona’ and ‘Tree view’ taxonomic classification visualizations available on TRAPID 2.0 are built using Krona (21) and the UniPept (22) visualizations (https://github.com/unipept/unipept-visualizations), respectively.

### Reference databases

Four reference databases are available within TRAPID 2.0: PLAZA 4.5 dicots and monocots (23), pico-PLAZA 3 (24), and eggNOG 4.5 (25). These databases integrate homology and functional information for genomes spanning a broad taxonomic range. Supplementary Table S2 provides an overview of their content and details the number of species, genes, and gene families they each incorporate.

In addition to these reference databases, leveraged to assign transcripts to gene families and functionally annotate them, TRAPID 2.0 makes use of the NCBI taxonomy (26) (downloaded 5^th^ of September 2019) for all taxonomy-related analyses.

### Similarity search, gene family assignment, and functional annotation

DIAMOND (27) was used to align query transcript sequences against a reference database, as a faster alternative to BLASTX. We run DIAMOND in ‘more sensitive’ mode (‘--more-sensitive’ flag), with a user-selected E-value cutoff set to 1e-5 by default. The proteome used as index for the similarity search can either be the full reference database or a user-selected taxonomy-constrained subset of it, corresponding to a specific phylogenetic clade or individual species. The latter will reduce the time needed for initial processing and would be suitable, for instance, to analyze data for which a closely related clade or species is available in the reference database.

For each transcript, the top protein similarity hit is retained and the gene family (GF) associated to this protein (if any) is assigned to the transcript. When using a PLAZA reference database, the functional annotation of each transcript can then either be transferred from the top protein hit from the similarity search, its assigned GF, or a combination of both (by default). This choice is also defined by the user. In case gene families are used to infer functional annotation, annotation labels (e.g. GO terms or proteins domains) that are transferred to the transcript have to be represented in at least 50% of the members of the GF (majority vote). To ensure the validity of this representation threshold, additional values were tested during functional annotation benchmark experiments conducted using data from five model species (see Supplementary Note S2).

When eggNOG 4.5 is used as reference database, the similarity search output file is processed using the annotation procedure of eggNOG-mapper (version 1) (28). For each transcript, the NOG associated to the seed ortholog (derived from the top protein hit of the similarity search) at the selected taxonomic scope is assigned as GF to the transcript. GO terms and KO terms reported by eggNOG-mapper are then transferred to the transcript. The taxonomic scope can be set by the user for all transcripts of an experiment, or adjusted automatically for each query (default).

Finally, if no protein hit was detected during the similarity search, no gene family or functional annotation are assigned to the transcript.

### Non-coding RNA identification

Potential non-coding RNAs present among query transcript sequences are identified using Infernal (29) with a selection of RNA models from Rfam 14.1 (30) (January 2019). By default, this selection comprises ubiquitous non-coding RNAs, with families from Rfam clans CL00001 (tRNA), CL00002 (RNase P), CL00003 (SRP RNA), CL00111 (small subunit rRNA), CL00112 (large subunit rRNA), and CL00113 (5S rRNA). Infernal’s ‘cmsearch’ command is run using parameters ‘--nohmmonly --rfam -- cut_ga’ (all models run in CM mode, heuristic filters set at Rfam-level, and Rfam’s gathering cutoffs used as reporting thresholds), as described in (31). The matched transcript sequences are assigned to their top Rfam RNA family (and clan, if any) and subsequently annotated with manually curated GO terms associated to the family. The mapping between Rfam RNA families and GO terms, defined by Rfam curators, was retrieved from Rfam’s FTP site (available at ftp://ftp.ebi.ac.uk/pub/databases/Rfam/14.1/database_files/database_link.txt.gz).

### Ribosomal RNA sequences post-processing

Transcripts assigned to any RNA family from CL00111 (SSU) and CL00112 (LSU) Rfam clans during TRAPID’s initial processing were considered as putative ribosomal RNA sequences. Their taxonomic classification was inferred using SINA (32) (version 1.5.0) with the SILVA 132 database (33) (using the ‘SSU Ref NR 99’ and ‘LSU Ref’ files with potential SSU and LSU sequences, respectively). SINA was set up to search and classify sequences using a minimum sequence similarity threshold of 0.9 and default values for all the other parameters.

### Assigning frame information, detection of potential frameshifts, and meta-annotation

For each transcript, retrieval of strand and frame information from similarity search hits (including detection of potential frameshifts) and homology-based ORF sequence prediction are performed using the strategy outlined in (18). In case the transcript does not have any similarity search hit and is not identified as being a non-coding RNA, its predicted ORF sequence is the longest ORF in all possible frames. The ORF sequence prediction step can use any of the genetic codes available from the NCBI taxonomy (https://www.ncbi.nlm.nih.gov/Taxonomy/Utils/wprintgc.cgi). It is additionally possible to run it after initial processing completion for sequences of individual transcript subsets, using any genetic code.

The meta-annotation of each transcript – a description of its full-length status – is based on a length comparison between its predicted coding sequence and the members of its associated GF, and the presence of predicted start and stop codons, as described in (18).

### Non-canonical genetic code use impact evaluation

Transcriptomes from 16 ciliate MMETSP samples (776,604 sequences in total) were processed with TRAPID 2.0, selecting eggNOG 4.5 as reference database, using either the standard genetic code (NCBI taxonomy translation table 1) or the ciliate nuclear genetic code (NCBI taxonomy translation table 6) for ORF prediction. All the other parameters were left at their default values. After initial processing, the ratio of predicted ORF sequence length over best similarity search hit length, termed ‘best hit recovery ratio’, was computed for all 257,454 transcript sequences having protein similarity search hits and the two genetic codes.

**Table 1.** Transcriptome annotation and analysis platform feature comparison. The abbreviations are as follows: KEGG Orthology groups (KO), Gene Ontology terms (GO), Protein Domains (PD), InterPro (IPR), BiGG metabolic reactions (BiGG), Enzyme Codes (EC), Conserved Domains Database (CDD), Differential Expression Analysis (DEA), and NCBI BLAST non-redundant database (NCBI NR).

| Features | Blast2GO (1) | KAAS | eggNOG-mapper v1 | Trinotate/Trinotate-web | EnTAP | Annocript | Dammit | TRAPID 2.0 |
| --- | --- | --- | --- | --- | --- | --- | --- | --- |
| Sequence similarity search | BLAST | BLAST | DIAMOND or HMMer | BLAST, HMMER | DIAMOND | BLAST, rpsBLAST | HMMER, LAST | DIAMOND |
| ORF finding | No | No | No | TransDecoder | GeneMarkS-T, TransDecoder | DNA2PEP | TransDecoder | Homology-supported/*de novo* using any NCBI translation table |
| Reference database | Public databases | Curated KEGG genes | eggNOG 4.5 | PFAM, Swiss-Prot | Up to 5 user-selected protein databases, eggNOG 4.5 | UniProt (or UniRef), CDD, TrEMBL, user protein databases | PFAM, OrthoDB, Uniref90, user protein databases | PLAZA 4 monocots/dicots, pico-PLAZA 2, eggNOG 4.5 |
| Gene family assignment | No | No | Yes | Yes | Yes | No | No | Yes |
| Functional ontologies | GO, PD (IPR), EC, KEGG | KO | GO, KO, BiGG, COG functional categories | GO, PD (PFAM, SignalP), KO | GO, PD (IPR), KO | GO, PD (PFAM) | GO, PD (PFAM) | GO, PD (IPR), KO |
| Non-coding RNA annotation | No | No | No | RNAMMER (rRNAs) | No | BLASTn with SILVA/Rfam, lncRNAs | Infernal with Rfam | Infernal with Rfam |
| Transcript expression use | No | No | No | DEA, clustering based on expression (RSEM) | Filtering of lowly expressed transcripts (RSEM) | DEA (Edge-R) | No | No |
| Functional enrichment analysis | Yes | No | No | Yes | No | No | No | Yes |
| Taxonomic classification | No | No | No | No | No | No | No | Kaiju with NCBI NR prot |
| Protein alignments | No | No | No (2) | No | No | No | No | MUSCLE, MAFFT |
| Phylogenetic trees | No | No | No (2) | No | No | No | No | FastTree2, IQ-TREE, RaxML, PhyML |
| Gene space completeness estimation | No | No | No | No | No | No | BUSCO | Core GF completeness |
| Interface | GUI | GUI | GUI and CLI | GUI and CLI | CLI | CLI | CLI | GUI |
| Unique features | Blast2GO Apps | Graphical pathway maps | Free text functional descriptions | Transmembrane regions (TMHMM) prediction | Similarity search filtering: Taxonomic Favoring and Contaminant Filtering | HTML summary (plots and statistics) | Ortholog detection using CRBL | Putative frameshift detection, ORF length meta-annotation, interactive analysis of transcript subsets |

(1) Blast2GO basic version
(2) MSAs and phylogenetic trees are available for each eggNOG group, externally of eggNOG-mapper

Protein sequences corresponding to two transcripts from MMETSP0018 (*Uronema sp. Bbcil*) that were assigned to ‘0IF5I’, an eggNOG orthologous group containing 100 proteins from 91 eukaryotes, were aligned with 4 reference sequences from Alveolata using MAFFT (34). Sequences predicted using the standard genetic code and the ciliate nuclear genetic code were used to compute the alignment. The MSA visual representation was generated using the ‘MSA’ R package (35) to create a TEXshade alignment (36).

### Estimation of gene space completeness using core gene families

TRAPID 2.0 estimates the gene space completeness of user-submitted transcriptomes by assessing the representation of core GFs. Core GFs are gene families that are highly conserved in a majority of species within a defined evolutionary lineage, and as such represent a valuable tool to estimate and analyze the gene space completeness in gene-based sequence data sets (37,38).

TRAPID 2.0’s core GF completeness analysis module enables the definition of a core GF set for any phylogenetic clade that is represented in the user-selected reference database, or any of the 107 available eggNOG taxonomic levels in the case of eggNOG 4.5, permitting the examination of gene space along an evolutionary gradient. Core GFs are defined using a simplified version of the approach described in (37): in order for a GF to be considered ‘core’ for a phylogenetic clade, it needs to be represented in at least 90% of the species of this clade. This threshold does not require complete conservation across all species of the clade and therefore tolerates potential annotation errors or GF loss in a limited number of species. It can alternatively be adjusted by the user in case more stringent or relaxed conservation requirements are needed. When defining a core GF set, each GF is given a weight based on its number of members and represented species as outlined in (37). The completeness analysis, performed after definition of the core GFs, consists of two steps. First, for each transcript, the top protein hit from the similarity search is retrieved, and the GF to which this protein is associated is considered to be represented. Then, a completeness score is computed (sum of the weight of represented core GFs on the weight of all core gene families) and reported, along with the list of represented and missing core GFs. In case the user chooses to use more than one top similarity search hit during the completeness analysis, gene families to which the top hit proteins are associated are scored based on the significance of the hits, and the GF having the best score is considered to be represented.

### Subset functional enrichment analysis

Subset functional enrichment analyses are performed using the hypergeometric distribution with a user-selected *q*-value cutoff ranging between 0.05 and 1e-5, and the annotation from all the transcripts of the experiment as background. For each enriched functional annotation label, the enrichment *q*-value is determined using the Benjamini–Hochberg correction for multiple hypothesis testing.

### Multiple sequence alignments and phylogenetic trees

Translated coding sequences of transcripts assigned to the same GF, combined with protein sequences of homologous genes from the reference database, are aligned using either MAFFT (34) or MUSCLE (39). The former uses automatic method detection with a maximum of 1,000 iterative refinement cycles while the latter uses default parameters. Input sequences used for the alignment can be filtered to by excluding reference protein sequences present in selected species and discarding transcript sequences flagged as ‘partial’ or arbitrarily selected. When creating a phylogenetic tree, optional editing of the alignment is performed using one of the available editing modes: removal of lowly conserved positions by trimming the sequences, filtering of partial sequences, or a combination of both. From this alignment, phylogenetic trees are inferred with either the approximate maximum likelihood program FastTree2 (40) or the maximum likelihood programs IQ-TREE (41), PhyML (42), or RaxML (43). Using IQ-TREE, trees are built under the best amino acid substitution model as selected by ModelFinder (44), chosen among a set of commonly used models (JTT, LG, WAG, Blosum62, VT, and Dayhoff). Rate heterogeneity across sites is accounted for using the FreeRate model (45), and branch support is estimated using the ultrafast bootstrap approximation (UFBoot) (46) with 1,000 bootstrap replicates. When using other programs, trees are built using the WAG substitution model in combination with 100 bootstraps and gamma approximation for modeling rate heterogeneity across sites. Generated Newick trees are converted to PhyloXML to enable incorporation of transcript meta-annotation and subset information. MSAs are visualized using the MSAviewer BioJS component (47,48) while phylogenetic trees are visualized using PhyD3 (49).

### Assembly, expression quantification, and analysis of diatom-dominated community metatranscriptomes

Raw pyrosequencing reads of samples obtained from diatom-rich communities of three sampling sites in the western Antarctic Peninsula (16) were retrieved from the NCBI Small Read Archive (Accessions SRX727358, SRX727361, SRX727362). Reads from each library were processed using Trimmomatic version 0.36 (50) to filter out short and low-quality sequences, and trim 5’ adapter sequences. Quality control of the sequences at various filtering stages was performed using FastQC version 0.11.2 (http://www.bioinformatics.babraham.ac.uk/projects/fastqc). Retained reads from all libraries (526,527 reads in total, see Supplementary Table S3) were combined to produce a ‘global’ metatranscriptome assembly, performed using MIRA version 4.0.2 (51). The resulting metatranscriptome consisted in 53,569 contigs, with a N50 contig length of 391 bp. A total of 208,592 reads were not used by MIRA to produce the assembly. Among these, 89,739 sequences longer than 200 bp after trimming were considered *bona fide* transcripts. After adding them to the metatranscriptome, a data set of 143,308 sequences (N50 320 bp) was obtained and used as input for further analyses.

Transcript expression quantification was performed as follows: first, filtered reads from each library were mapped to the metatranscriptome (contigs and retained singletons) with BWA mem (52) version 0.7.17. Second, read counts were computed using FeatureCounts (subread package version 1.6.2) (53). Finally, read counts were transformed into TPM values and used as a basis for transcript subset definition. For each sampling site, subsets containing all expressed transcripts were defined, considering transcripts having a TPM value of 2 or more for a given library to be expressed in the corresponding sampling site and filtering out identified non-coding RNAs.

Transcript sequences were processed using TRAPID 2.0, selecting pico-PLAZA 3 as a reference database. All the initial processing parameters were set to their default values (‘Eukaryotes’ as clade and 1e-5 maximum E-value cutoff for the similarity search, functional annotation transfer from both GF and best similarity search hit, default Rfam clans for Infernal, and taxonomic classification enabled). All the transcript subsets were subsequently loaded into TRAPID 2.0 and refined to only retain sequences assigned to diatom (i.e. assigned to ‘Bacillariophyta’) to create diatom-specific subsets expressed in each sampling site, and subset functional enrichment analyses were performed from the web application to examine the functional variations between the three phytoplankton communities.

### Implementation

The TRAPID 2.0 web application was implemented using CakePHP (https://cakephp.org) and a MySQL database (https://www.mysql.com) as back-end. The available interactive visualizations were developed using two open-source JavaScript libraries, D3.js (https://d3js.org) and Highcharts (https://www.highcharts.com), or built on top of other open-source software projects mentioned in the above sections. TRAPID’s analysis pipelines consist of a collection of programs and scripts written in Java, Perl, and python. All TRAPID 2.0 jobs run on a computing cluster (64 Intel(R) Xeon(R) Platinum 8153 cores, 264 GB of memory). Information regarding the status of jobs and the load of the cluster is available via the job management page.

## RESULTS AND DISCUSSION

### A two-phase approach for the annotation and exploration of assembled transcriptome data

TRAPID 2.0 is a web application for the annotation and exploration of assembled transcriptome data. It processes a set of input sequences, named an experiment, in two distinct phases: the initial processing and the exploratory phases (Figure 1). The individual steps they incorporate (Supplementary Table S4) tackle diverse aspects of the characterization of transcriptomes and enable the end-user to address a variety of biological questions, including transcript quality control, the assignment of transcripts to gene or RNA families, and the generation of functional annotations.

**Figure 1.**
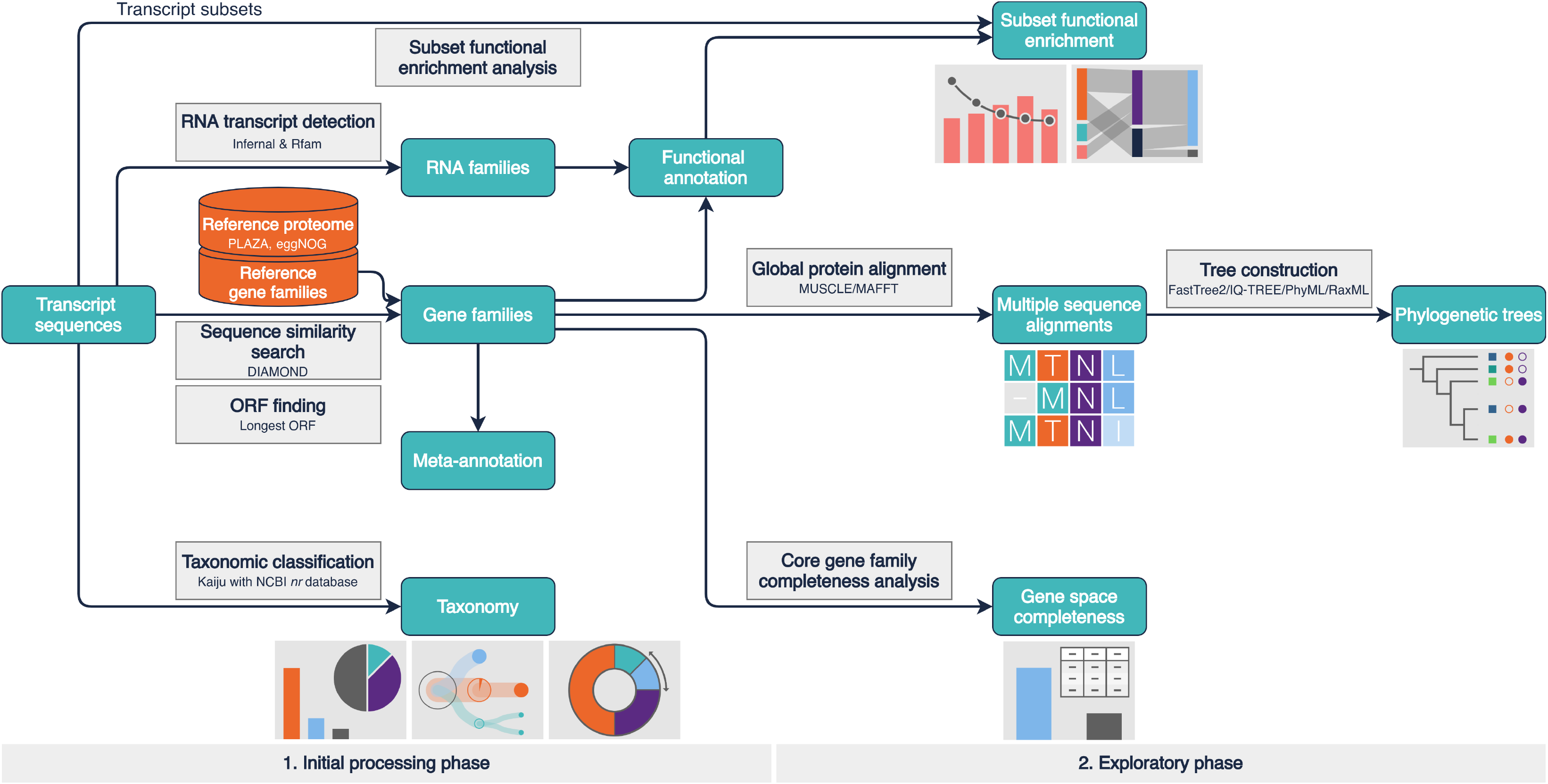
TRAPID 2.0 workflow and functionality overview. TRAPID’s workflow comprises two main parts: an initial processing phase (1), executed non-interactively after data upload, and an exploratory phase that consists of a set of functional and comparative tools (2) accessible through the web application. Input and output data are represented by cyan boxes. Output data boxes are complemented by thumbnails that depict the visualizations available from the web application. Available reference databases (consisting of functionally annotated proteomes and gene families) are represented by orange cylinders. Analysis and computation steps are represented by grey boxes and solid arrows.

Executed non-interactively, the initial processing phase ingests, analyzes, and extracts meaningful information from the input data. The first step, the taxonomic classification of transcripts, enables the detection of potential contaminant sequences and facilitates the characterization and analysis of metatranscriptomes. It is followed by an RNA homology search to identify putative non-coding transcripts, subsequently assigned to RNA families and functionally annotated. Next, TRAPID 2.0 leverages predefined reference databases, which contain biological sequences from multiple species clustered in precomputed homologous gene families (GFs) and having functional annotations. A sequence similarity search against a proteome from one of the available reference databases is employed to assign transcripts to GFs, transfer homology-based functional annotations, infer coding sequences, identify putative frameshift errors, and estimate the full-length status of each transcript. In case the user chooses to process untranslated coding sequences rather than transcripts, for instance predicted from a genome or resulting from any other sequencing effort, the ORF finding routine is skipped: all sequences are translated using frame and strand ‘+1’ with the user-selected genetic code.

The exploratory phase features a collection of components accessible via the web interface for downstream analysis and detailed data inspection, exploiting the previously generated information. The experiment statistics report comprehensive annotation metrics and allow the examination of the length distribution of transcripts and predicted ORF sequences. The core GF completeness module provides a means to estimate gene space completeness at varying evolutionary levels by assessing the representation of conserved, quasi-universal gene families within the input data. After defining transcript subsets, functional biases within subsets can be identified and investigated through subset functional enrichment analysis and interactive gene function visualization tools. Transcript subsets can additionally be compared to identify their unique or shared functional characteristics. Finally, transcript sequences can be studied alongside their detected homologs in an evolutionary context to identify orthologs and paralogs, using multiple sequence alignments and phylogenetic trees. Data processing and analyses performed within TRAPID 2.0 are transparent and reproducible: the experiment log stores extensive information about all computation steps, applied tools, and parameters.

### Taxonomic classification of transcript sequences to detect putative contaminants and disentangle microbial communities

After the user creates a TRAPID experiment and uploads assembled transcripts (or coding sequences), they are taxonomically classified. This initial processing step serves a dual purpose: it enables the identification of potential contaminant sequences within single-species transcriptomes, and facilitates the analysis of transcriptome data from communities. It may be disabled by the user in case it is considered unrelated to the analysis at hand. The taxonomic classification is performed using Kaiju (20), a program based on finding maximal exact matching substrings between queries and protein sequences of a reference database. To enable the classification of transcripts from the broadest possible taxonomic range and to remain as unbiased as possible with regards to the selected clades used as reference, the database used by Kaiju consists of 219 million sequences from eukaryotes, bacteria, archaea, and viruses in the NCBI non-redundant protein database. Although several taxonomic classification methods have been described, see e.g. (54–56), Kaiju was ultimately retained for this step due to its low running time, its ability to overcome large evolutionary divergences, and because it was shown to be more sensitive than *k-mer*-based methods to classify sequences from clades that are underrepresented in reference databases (20), making it an appropriate candidate for the study of non-model species. Kaiju assigns transcript sequences to a species or strain, or to a higher-level node in the taxonomic tree in case of ambiguity: if a transcript contains a protein fragment that is identical between two organisms from the same clade, then it is assigned to this clade, which represents the lowest common ancestor of these organisms (20). Upon completion of the initial processing, taxonomic classification results are made available to the user as multiple interactive visualizations, together with a tab-delimited file containing raw results (Figure 2). The Krona sunburst chart (Figure 2A) and the tree viewer (Figure 2B) enable an in-depth examination of the results through the exploration of the complete taxonomic tree, whereas the sample composition bar and pie charts (Figure 2C) provide a quick overview of the results, depicting the domain-level composition and the ten most represented clades at adjustable taxonomic ranks. Any of these interactive visualizations can also be used to rapidly create transcript subsets by selecting clades: all the transcripts assigned to the selected clades are then grouped in a new transcript subset that can be further analyzed within TRAPID 2.0.

**Figure 2.**
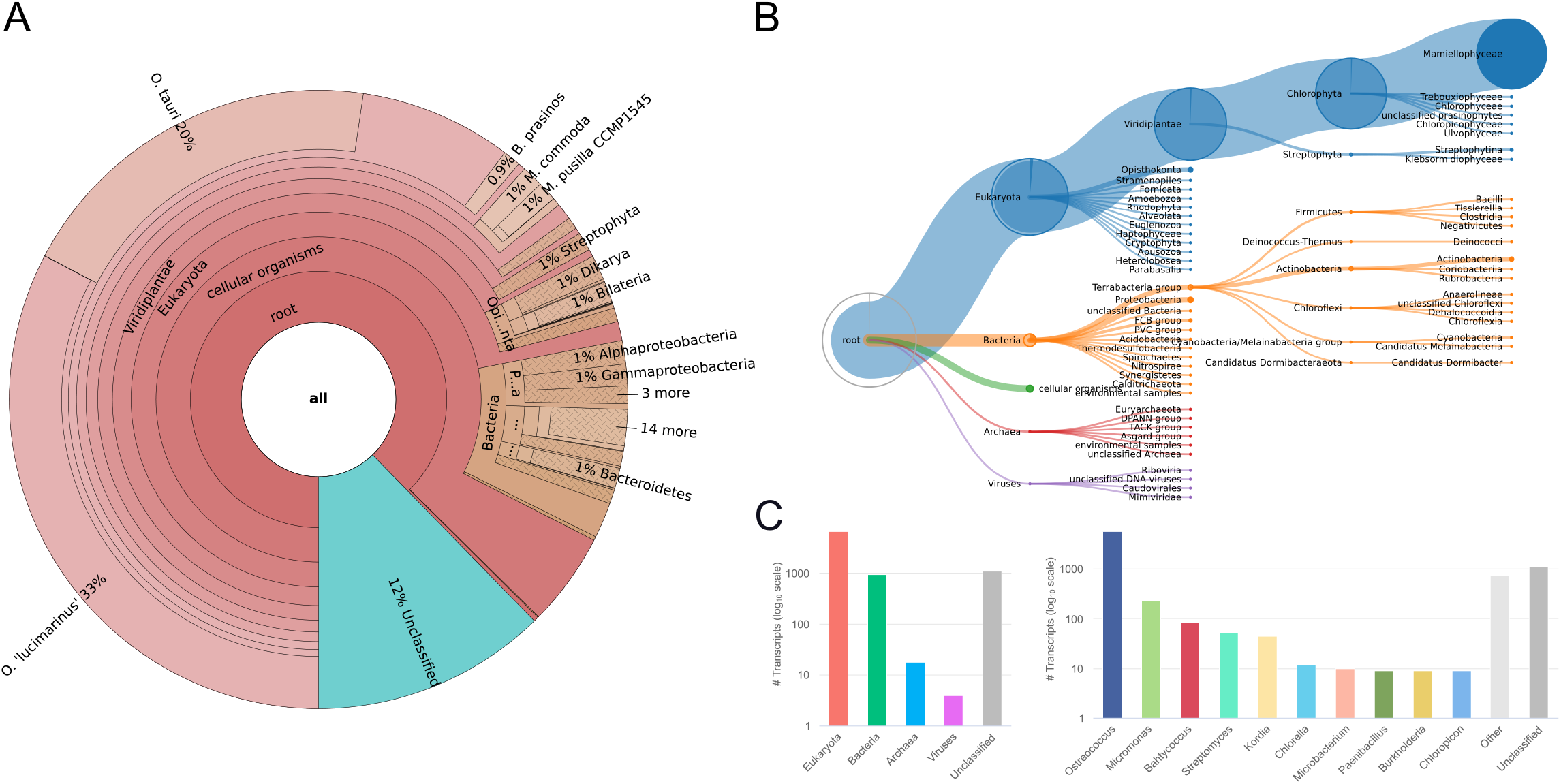
*Ostreococcus mediterraneus* transcriptome taxonomic classification results visualization. (**A**) Krona sunburst chart. (**B**) Tree viewer: users can explore the classification of transcripts along the tree of life, enabling in-depth investigation of the results. (**C**) Sample composition bar charts: domain-level composition is shown on the left and the top ten most represented clades at a given taxonomic rank (here genus) on the right, providing an overview of the results. Transcript subsets can be defined using any of the available visualizations for further analysis. These results were generated using MMETSP0929 (*Ostreococcus mediterraneus* clade-D-RCC2572).

Benchmark experiments were performed to assess Kaiju’s performance with regards to the classification of assembled transcript sequences and to explore various running mode and parameter combinations (Supplementary Note S1; Supplementary Figure S1). Based on the results obtained after evaluating Kaiju with two types of evaluation data sets having different properties, we concluded the selected running mode combines good genus-level classification performance with computational efficiency.

Genus-level taxonomic classification results for 18 microbial eukaryote transcriptomes from the Marine Microbial Eukaryote Transcriptome Sequencing Project (MMETSP) (9) provides insights into their quality and ecology (Figure 3A). While overall large fractions (21.9- 89.0%) of transcripts were assigned to eukaryotic genera matching sample metadata, nine different bacterial genera were present among the most represented genera for the 18 inspected samples. This observation likely reflects the general low level of contamination problem mentioned in the MMETSP publication, with contaminant sequences originating from species living in culture with the sequenced organism (57,58) or from experimental procedures (e.g. during library construction and sequencing). However, the four analyzed diatom transcriptomes (‘MMETSP0737-740’), all corresponding to *Thalassosira miniscula* samples, exhibited varying fractions of transcripts assigned to bacteria (mostly Rhodobacterales), with up to 30.2% assigned transcripts for MMETSP0740 in our summary, suggesting for these samples contamination levels may have been underestimated. Three samples of the labyrinthulid hard clam parasite QPX (‘MMETSP0098-100’), obtained from clam tissue, displayed similar phylum-level taxonomic classification profile, with a large fraction of unclassified sequences (up to 54.6% from ‘MMETSP0100’) and a high diversity of represented phyla. Interestingly, for MMETSP0098, 26.4% of transcripts were assigned to Crassostrea, potentially representing sequences from the host. In general, the observed unclassified fractions point to the lack of available reference genomic information for microbial eukaryotes, motivating further sequencing efforts (59), or to the presence of misassembled transcripts or untranslated sequences lacking protein similarity to known genes.

**Figure 3.**
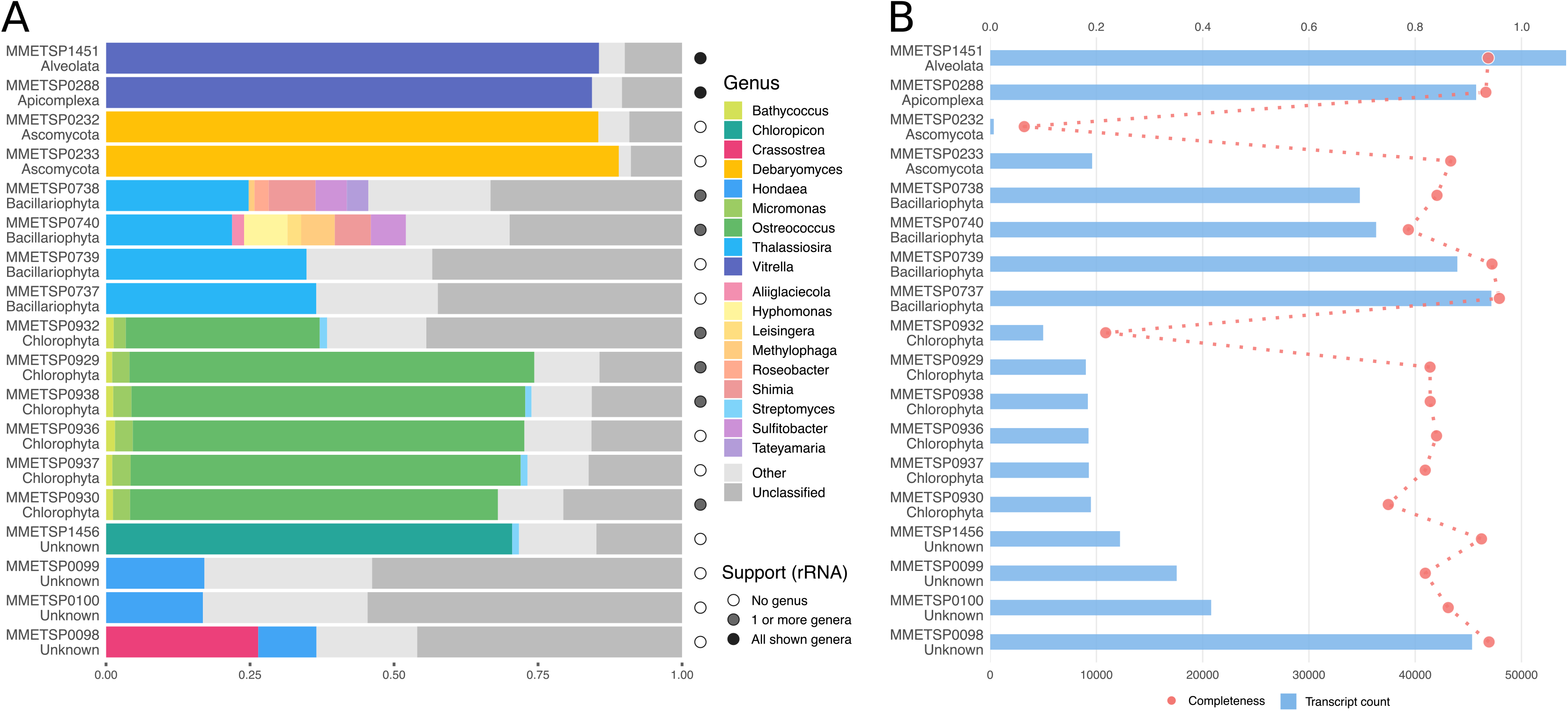
Genus-level taxonomic classification, core eukaryotic completeness, and transcript count of 18 microbial eukaryote transcriptomes. (**A**) Genus-level taxonomic classification summary. For each transcriptome, only up to 10 top represented genera (genera to which the most transcripts assigned to) are shown. Transcripts assigned to other genera or genera encompassing less than 1% of the sample are aggregated as ‘Other’ (light grey fraction), transcripts not assigned to any taxonomic label are represented as ‘Unclassified’ (dark grey fraction). The circle on the right of each classification summary indicates the number of top represented genera additionally supported by the classification of either SSU or LSU rRNA sequences detected by TRAPID 2.0 (see Methods). (**B**) core eukaryotic completeness score, computed using 1,116 ‘core’ eggNOG orthologous groups conserved in at least 90% of the eukaryote organisms present in the database (red dots, top x-axis), and transcript count (blue bars, bottom x-axis). The phylum label associated to each depicted transcriptome was retrieved from the MMETSP metadata. These results were generated using 18 MMETSP samples processed with eggNOG 4.5 as a reference database and default initial processing parameters.

Apart from the characterization of protein-coding transcripts, TRAPID 2.0 identifies potential non-coding RNAs using Infernal (29) with a selection of RNA models from Rfam (30), representing ubiquitous non-coding RNAs by default (see Material and Methods). Identified non-coding transcripts are assigned to the corresponding Rfam family and clan, if any, and are subsequently annotated with *associated* GO terms. Identified RNA families can be browsed (Supplementary Figure S2), Infernal’s output examined via individual transcript pages, and RNA transcripts easily exported for further characterization, for example using the SILVA database (33) to examine/query predicted ribosomal RNA sequences or any other dedicated tool, such as tRNAscan-SE to annotate tRNAs (60).

Comparing the genus-level classification results of Kaiju, based on all the transcripts of each sample, with the classification of SSU/LSU ribosomal RNAs identified by TRAPID 2.0 revealed that in 44% (8/18) of the samples the most abundant genera were supported by ribosomal RNA analysis (Figure 2). For only two samples (MMETSP1451 and MMETSP0288, the two first samples in Figure 3A) all most abundant genera were recovered, demonstrating the complementarity of protein-based and ribosomal RNA based taxonomic classification approaches.

### Homology-based functional annotation

After completion of the taxonomic classification, DIAMOND (27) is employed to perform a sequence similarity search against a user-selected protein database (Figure 1). The used protein database consists of protein sequences retrieved from one of the available reference databases, which contain functionally annotated biological sequences from multiple species clustered in precomputed homologous gene families (Supplementary Table S2). By default, the entire reference proteome (proteins from all the species of the reference database) is used, but it can optionally be constrained to a subset of proteins originating from a specific phylogenetic clade or individual species.

Within TRAPID 2.0, four reference databases are available: PLAZA 4.5 dicots and monocots (23), pico-PLAZA 3 (24), and eggNOG 4.5 (25). Taken together, they encompass protein sequences, functional annotation, and GF information for 115 archaea, 1,678 bacteria, and 319 eukaryotes (81 of which exclusively present in PLAZA databases). These reference databases span a broad and diverse taxonomic range, providing flexibility to the user and making it possible to process transcript sequences of diverse origin. They furthermore represent a high-quality backbone for the comparative and functional analyses that can subsequently be performed during the exploratory phase.

The similarity search output is utilized during a post-processing step to provide input sequences with several layers of information (Supplementary Table S4). First, each transcript is assigned to a gene family from the reference database, and this information is used to transfer functional annotation to the transcript (GO terms, InterPro domains, or KO terms). Second, using the alignments reported for each transcript, frame statistics are generated and used to infer the transcript’s ORF sequence, i.e. the longest ORF within the frame supported by sequence similarity hits, and to predict whether the transcript contains putative frameshifts in case conflicting frame information was detected. Finally, meta-annotation, an estimation of the full-length status of the transcript, is generated based on the comparison of the predicted ORF sequence length to the coding sequence length of the genes from the GF the transcript was assigned to. In case eggNOG 4.5 was selected as reference database, the GF assignment and functional annotation transfers are performed using the eggNOG-mapper annotation routine (28).

To evaluate the performance of the functional annotation procedure, we compared the GO annotations inferred by TRAPID 2.0 to gold standard GO annotations for cDNA sequences originating from five well-characterized model organisms (*Arabidopsis thaliana, Drosophila melanogaster, Escherichia coli (strain K12), Homo sapiens,* and *Saccharomyces cerevisiae;* see Supplementary Note S2). For each organism and GO aspect, recall, precision, F_1_ score, and annotation coverage were determined when either eggNOG 4.5 or PLAZA 4.5 dicots was selected as a reference database, additionally evaluating multiple GF transfer threshold values for the latter (Supplementary Figure S3; Supplementary Note S2). Due to the broad taxonomic distribution of species in eggNOG 4.5, encompassing all the evaluated organisms, a better overall functional annotation performance (higher F_1_ score) was achieved when it was used as a reference database compared to PLAZA 4.5 dicots, with the exception of *Arabidopsis thaliana*. Considering that all the evaluated organisms are represented within eggNOG 4.5, the observed annotation coverage was in some cases low (e.g. ranging between 24.2 and 66.1% for *Escherichia coli*), potentially linked to the fact that only non-electronic annotations are transferred by eggNOG-mapper, and because not all proteins necessarily have associated GO annotations. Finally, since *Arabidopsis thaliana* is represented in both PLAZA 4.5 dicots and eggNOG 4.5, focusing on the evaluation results for this species enabled us to compare the functional annotations inferred using both reference databases. Processing *Arabidopsis thaliana* cDNA sequences using PLAZA 4.5 as a reference database yielded better annotation metrics compared to using eggNOG 4.5 (higher F_1_ score for the ‘biological process’ and ‘molecular function’ GO aspects and higher annotation coverage for all GO aspects). Drawing on this evaluation, we recommend users to select one of the available PLAZA instances as a reference database in case they annotate transcript sequences originating from plants or microbial photosynthetic eukaryotes. However, eggNOG 4.5 remains a suitable default choice, due to the broad taxonomic distribution of represented species, its taxonomy-constrained orthologous groups, and the functional annotation performance achieved by eggNOG-mapper.

### ORF finding using non-canonical genetic codes

The genetic code, the translation of nucleotide triplets into amino acids, is universal in nearly all organisms and cellular locations. While it was termed a ‘frozen accident’ (61) and long considered to be completely unalterable, variations of the standard genetic code have been observed within the tree of life as well as in organellar genomes (62). Among others, notable examples of non-canonical genetic code use reported for eukaryotes include the reassignment of stop codons by certain ciliates (63,64) and ulvophycean green algae (65).

To accommodate these variations, TRAPID 2.0 supports the use of non-canonical genetic codes during ORF detection and translation with any of the genetic codes available from the NCBI taxonomy. In practice, the user may select a non-canonical genetic code to use for all the transcripts of an experiment prior to initial processing. After completion of the initial processing, it becomes possible to perform ORF finding and translation using any genetic code for individual transcript subsets, based for instance on taxonomic classification results, via the subset page.

To evaluate the impact of using an appropriate genetic code during ORF finding, 16 ciliate MMETSP transcriptomes (825,773 sequences) were processed with TRAPID 2.0 selecting either the standard or the ciliate nuclear genetic code for ORF detection (see Material and Methods for individual accessions and initial processing parameters). Transcript sequences having a protein similarity search hit against the reference database were retained and the length of their predicted ORF sequence was compared with the length of their best hit (Figure 4A). Performing the initial processing using the ciliate nuclear genetic code yielded ORF sequences that better support similarity search results, longer and closer to the best hit length, compared to the sequences predicted using the standard genetic code. The quantity of transcripts flagged as ‘partial’ moreover decreased by 28.4% (from 166,011 to 118,815), indicating the recovery of more complete protein sequences. To illustrate how downstream, GF-level analyses are impacted by the use of non-canonical genetic code, we aligned two *Uronema* transcripts with four Alveolata reference sequences from their assigned eggNOG orthologous group, using proteins predicted with both the standard or the ciliate nuclear genetic code (Figure 4B). Longer amino acid sequences having higher global similarity were recovered when the ciliate nuclear genetic code was used during initial processing (stop codon reassignment), whereas substantially shorter amino acid fragments were recovered when using the standard nuclear genetic code.

**Figure 4.**
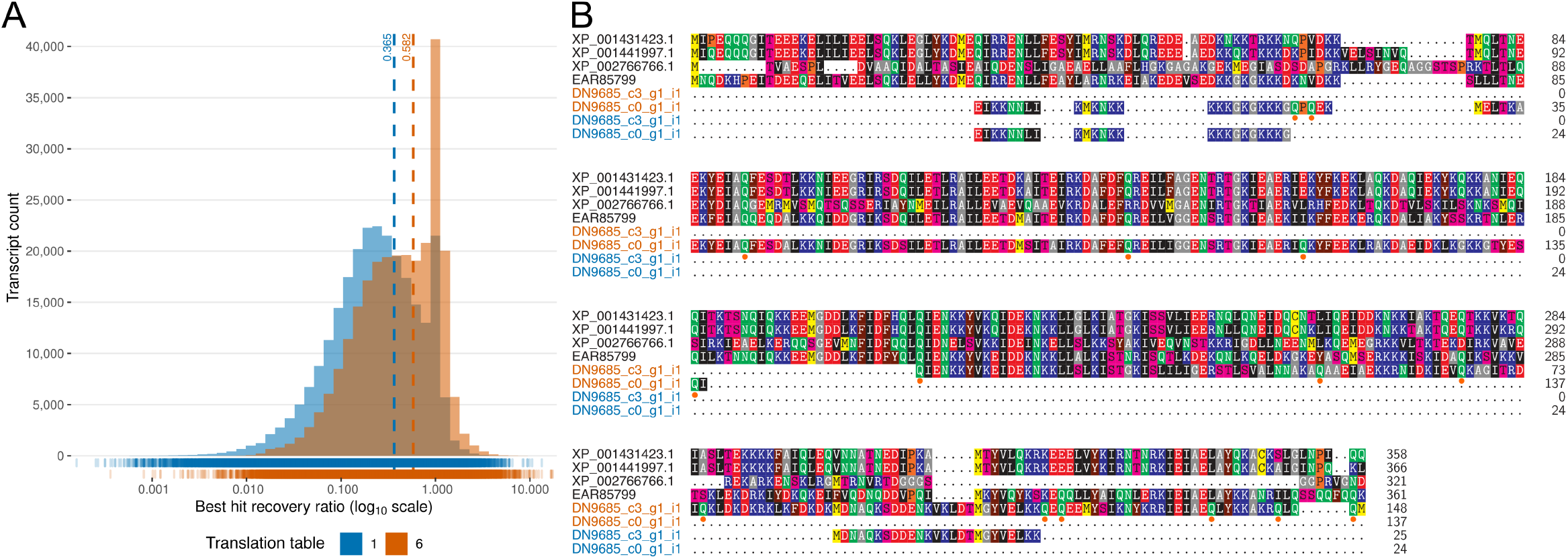
Impact of non-canonical genetic code use during ORF prediction for ciliate transcriptomes. (**A**) Histogram of predicted ORF sequence length divided by best sequence similarity search hit length (‘best hit recovery ratio’) for 257,454 sequences from 16 ciliate MMETSP samples having homology support, using the standard or the ciliate nuclear genetic code for ORF prediction (translation table 1 or 6 respectively). For each genetic code, the distribution of ratio values is depicted as a rug plot, and the mean value represented by a dashed line. (**B**) Multiple sequence alignment of 2 transcripts from MMETSP0018 (*Uronema sp. Bbcil*) assigned to the ‘0IF5I’ eggNOG orthologous group with reference sequences from Alveolata. ORF sequences were predicted using either the standard or the ciliate nuclear genetic code (corresponding to blue and orange sequence labels, respectively). Amino acid residues are shaded based on the chemical properties of their functional groups, with orange circles indicating stop codons reassigned to glutamine. Sequence label prefixes were trimmed to improve legibility.

### Rapid estimation of gene space completeness along an evolutionary gradient

The thorough evaluation of assembled transcriptomes is critical, as downstream analyses are directly impacted by quality. While technical sequencing quality metrics enable to estimate assembly contiguity (e.g. metrics returned by TransRate (66)), the information they provide only partially reflects the quality of the data and they should therefore be complemented by metrics that evaluate the global gene content. Such metrics frequently compare the observed and the expected gene content, the latter being modeled using a set of evolutionarily conserved marker genes that can range from ancestral, highly conserved genes, to clade- or species-specific genes (38). TRAPID 2.0 leverages the GF assignment step of the initial processing, enabling users to assess and analyze the gene space completeness of transcriptomes (Supplementary Figure S4) by inspecting the representation of core gene families (‘core GFs’). Core GFs consist of a set of gene families that are highly conserved in a majority of species within a defined evolutionary lineage (more details in Material and Methods). A key feature of this functionality in TRAPID 2.0 is the on-the-fly definition of core GF sets for any lineage represented in the selected reference database, making it possible for users to rapidly examine gene space completeness along an evolutionary gradient and check if core/ancestral conserved genes or clade-specific genes are represented within their data set (38). The main output of the core GF completeness analysis is the completeness score, an intuitive quantitative measure of the gene space completeness at the selected taxonomic level ranging between 0-100%. The represented or missing core GFs and their associated biological functions are also reported, enabling the identification of potential functional biases.

The possibility to define core GF sets for any lineage is one of the two major differences between our approach and BUSCO (67), a widely used tool for gene space completeness evaluation based on the presence of quasi single-copy genes predefined for major clades (68). The second difference is the use of multi-copy gene families. The usage of all conserved gene families, regardless of their copy number, enables to cover a larger fraction of the gene function space and reduces strong functional biases. For instance, many conserved genes in eukaryotes are not single-copy, such as histones, transcriptions factors, or other key components of the eukaryotic cell machinery (69).

Figure 3B shows an overview of transcript counts and eukaryotic gene space completeness, computed using 1,116 core eukaryotic eggNOG orthologous groups, for 18 microbial eukaryote samples from the MMETSP. Overall, the observed transcriptomes exhibit high core eukaryotic GF completeness scores, apart from ‘MMETSP0932’ and ‘MMETSP0232’. With the exception of transcriptomes containing a very small number of sequences (e.g. MMETSP0232 with 342 sequences), number of transcripts and core eukaryotic completeness score are not strongly correlated. For the analyzed Bacillariophyta samples (all corresponding to *Thalassiosira miniscula*), the variation in the fraction of transcripts assigned to bacterial phyla matches the variation in completeness scores, revealing the high occurrence of Proteobacteria in two of these diatom samples (MMETSP0740 and MMETSP0738). Finally, the presence of large fractions of unclassified sequences does not seem to result in lower completeness scores, illustrating that these two measures capture different information and the use of a combination of evaluation metrics should be employed to assess the quality of transcriptomes.

### Functional analysis and comparison of transcript subsets

In addition to the global characterization of transcriptomes, the detailed analysis and comparison of transcript subsets can lead to valuable biological insights. Starting from any arbitrary list of transcript identifiers (for instance, transcripts expressed in a specific condition), TRAPID 2.0 users can define subsets and use them to perform subsequent functional analyses. A new transcript subset may either be uploaded as a file or created interactively from the web application, for example based on the taxonomic classification results or the annotation as protein-coding or RNA gene. Three default subsets are defined upon initial processing completion, encompassing sequences that were assigned to a gene family (protein-coding transcripts), an RNA family (RNA transcripts), and both a gene and an RNA family, (ambiguous transcripts, potentially marking misassembled or fused transcripts).

Individual transcript subsets are characterized through functional enrichment analysis, using the annotation from all the transcripts of the experiment as background. Functional enrichment results are reported as a bar chart depicting the enrichment fold and *q*-value (corrected *p*-value) of significantly enriched functional annotation labels, as shown in Figure 5A for 48 *Ostreococcus mediterraneus* metal ion transport transcripts (MMETSP0936). In the case of GO enrichment analysis, a subgraph representing the hierarchy of enriched GO terms within each aspect can also be viewed (Supplementary Figure S5). The complex relationship between transcript subsets, enriched functional annotation labels, and associated gene families can additionally be investigated using interactive Sankey diagrams (Figure 5B).

**Figure 5.**
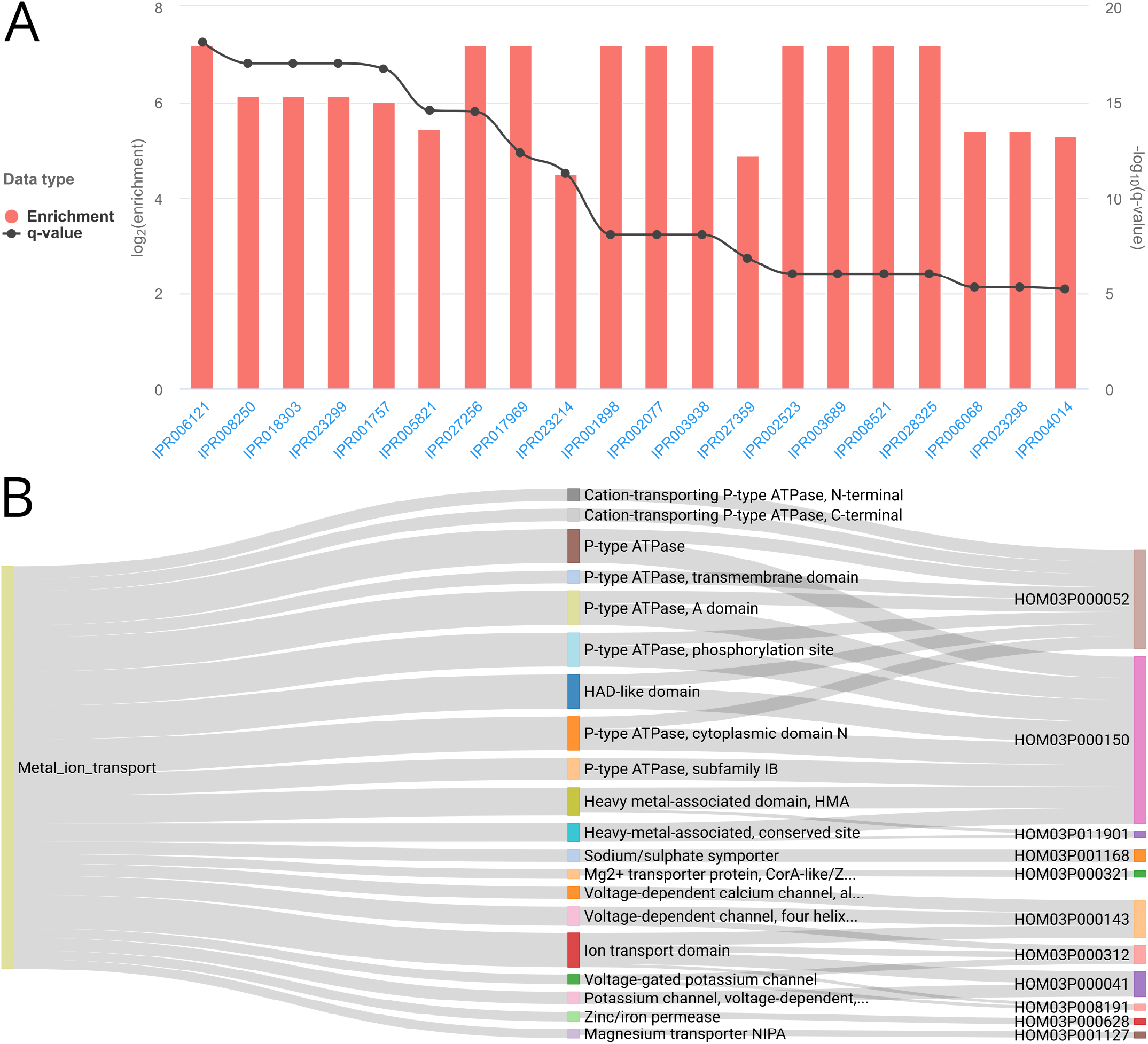
Functional enrichment analysis of a subset of *Ostreococcus mediterraneus* metal ion transport transcripts. (**A**) Metal ion transport transcripts InterPro enrichment results. InterPro identifiers are represented on the x-axis, enrichment fold on the left y-axis (red bars), and enrichment *q*-value on the right y-axis (dark grey dots). (**B**) Sankey diagram depicting the relationships between metal transport transcripts (left blocks), significantly enriched InterPro domains (middle blocks), and PLAZA gene families (right blocks). Line width is proportional to transcript annotation (left lines) and GF membership (right lines). The maximum enrichment *q*-value threshold is 1e-5 and only gene families containing at least two transcripts are displayed. These results were generated using MMETSP0936 (*Ostreococcus mediterraneus* clade-D-RCC2573) with pico-PLAZA 3 as a reference database and default initial processing parameters.

Besides the analysis of individual subsets, TRAPID 2.0 supports their comparison. By computing functional annotation ratios between subsets, it is possible for the user to identify shared and unique functions. The interactive Sankey diagrams presented in Figure 5B also support the examination of multiple subsets simultaneously, given functional enrichment analysis was performed for them.

### Comparison with other transcriptome analysis tools

While numerous transcriptome analysis tools are publicly available, a comparison of their functionalities highlights TRAPID 2.0’s unique properties (Table 1). We selected seven tools for comparison based on their similarity with TRAPID 2.0 and community usage: Blast2GO (70), KAAS (71), eggNOG-mapper (28), Trinotate (coupled with Trinotate-web) (72), EnTAP (73), Annocript (74), and dammit (https://doi.org/10.5281/zenodo.3569831). Since several of these tools are web services or rely on web services, a comparison of their execution time would likely be biased as it may be influenced by variables that cannot be controlled for, such as the underlying computational resources or the current server load. For the end-user, the time required to process a transcriptome nevertheless remains a relevant consideration in choosing a tool. Therefore, we report TRAPID 2.0’s initial processing execution times for five different microbial eukaryote transcriptomes and all available reference databases (Supplementary Figure S6). As one would intuitively expect, the execution time depended heavily on the size of the input data set and the used reference database. TRAPID 2.0 processed the largest tested transcriptome (119,699 sequences) in under five hours in all settings.

Although all the compared tools perform sequence similarity search against reference proteins and functional annotation of input sequences, these two tasks are performed using different methods, reference databases, and functional ontologies, resulting in divergent speed, accuracy, and comprehensiveness. Other features are shared by a limited number of tools exclusively. For instance, only eggNOG-mapper, Trinotate, EnTAP, and TRAPID 2.0 assign input sequences to gene families. In addition, only Trinotate, EnTAP, Annocript, dammit, and TRAPID 2.0 include the prediction of ORF sequences as a processing step, since they were developed for the analysis of *de novo* transcriptomes in particular. Moreover, while half of compared tools focus on the annotation of protein-coding sequences, only Trinotate, Annocript, dammit, and TRAPID 2.0 take non-coding RNAs into account. In terms of accessibility, eggNOG-mapper, Blast2GO, KAAS, and TRAPID 2.0 feature user-friendly graphical interfaces for the processing of input sequences, easing their adoption by a wider community than command-line tools. The interfaces of TRAPID 2.0, Blast2GO, and Trinotate web (accessed via a local webserver) support subsequent data exploration, after sequences were processed. Additionally, TRAPID 2.0 offers some unique features including the detection of putative frameshifts and built-in taxonomic classification.

Besides the multilayered annotation that TRAPID 2.0 provides during its initial processing phase, comprising structural, functional, and taxonomic information, the main difference between TRAPID 2.0 and the other examined tools resides in its exploratory phase. In addition to permitting an in-depth exploration of the generated results, TRAPID’s analytical toolkit enables users to address a wide array of biological questions directly from the web application. Using an intuitive and interactive interface, it is possible to inspect sequence conservation and evolution, estimate gene space completeness (also possible via command-line with dammit using BUSCO), and perform functional enrichment analyses and detailed comparisons of subsets.

Finally, in contrast to Trinotate, EnTAP, and Annocript, TRAPID 2.0 does not perform transcript expression quantification nor provides any straightforward way to directly incorporate this information. It is nevertheless possible to perform expression quantitation externally and define transcript subsets based on observed expression patterns to conduct downstream analyses via TRAPID 2.0, as illustrated in the next section.

### Studying taxonomic and functional variation in diatom community metatranscriptomes

To demonstrate TRAPID’s capabilities, we employed it to analyze metatranscriptomics data of three samples obtained from diatom-dominated phytoplankton communities (16). The samples were collected from three distinct sites in the western Antarctica Peninsula: the Bransfield Strait (BFS; 30 m), the Weddell Sea (WDS; 6-45 m), and the Wilkins Ice Shelf (WKI; melted sea ice). After quality control and filtering, 526,527 pyrosequencing reads from all samples were assembled into 53,569 contigs, with a N50 contig length of 391 bp (Supplementary Table S3). Singletons longer than 200 bp after trimming, considered *bona fide* transcripts, were subsequently incorporated into the metatranscriptome, resulting in a data set of 143,308 transcript sequences (N50 320 bp) used as input for TRAPID 2.0.

To characterize and compare those phytoplankton communities, we first quantified expression in each sampling site (Supplementary Table S3). After generating and transforming raw counts, multiple transcript subsets were defined based on their expression patterns across the three sampling sites: for each site, subsets containing all the expressed transcripts (TPM >= 2) were defined (see Material and Methods), filtering out identified non-coding RNAs.

Transcript sequences were subsequently processed with TRAPID 2.0, using pico-PLAZA 3 as reference database and default parameters (see Material and Methods). 63,641 (44.4%) transcripts were assigned to 8,931 gene families and 4,536 transcripts (3.2%) to 13 RNA families. 77,105 (53.8%) sequences received a taxonomic classification. 65,831 (45.9%) transcripts were functionally annotated with 12,965 distinct GO terms, and 63,641 (44.4%) transcripts with 7,052 distinct InterPro domains (Supplementary Table S5). Following uploading of the defined transcript subsets into TRAPID 2.0, we refined the subsets to only include sequences assigned to Bacillariophyta and performed subset functional enrichment analyses from the web application to examine the functional variations between the three diatom communities.

A domain-level overview of the taxonomic classification of the protein-coding transcripts of the global metatranscriptome reveals that Eukaryota is the most represented domain (Figure 6A), the remainder of transcripts being mainly assigned to Bacteria, with only less than 1% assigned to Archaea or viruses. At the genus-level, the inspection of the taxonomic classification of transcripts expressed in each sampling site shows a global domination by diatom transcripts in the three sites and site-specific variations (Figure 6B). Transcript sequences were mainly assigned to *Fragilariopsis*, *Pseudo-nitzschia*, and *Thalassiosira*, in contrasting proportions depending on the sampling site. The BFS sample features the largest fraction of transcripts assigned to *Pseudo-nitzschia* (20% of sequences assigned to eukaryotes and 33% of sequences assigned to diatoms), WDS the highest fraction assigned to *Thalassiosira* (23% of sequences assigned to eukaryotes and 52% of sequences assigned to diatoms), and WKI the highest fraction assigned to *Fragilariopsis* (23% of sequences assigned to eukaryotes and 48% of sequences assigned to diatoms). The observed repartition of protein-coding transcripts assigned to diatom genera per sampling site supports results previously obtained with a ribosomal protein gene maximum likelihood phylogenetic analysis (14). Interestingly, WKI exhibits a substantial portion of transcripts assigned to *Eurytemora* and *Tigriopus* (5% of sequences assigned to eukaryotes), two genera of calanoid copepods. The presence of copepods in this environment is not unusual, as they represent a key link between primary production and higher trophic modes, grazing on phytoplanktons. One such example is *S. longipes*, a species observed in a similar environment (location and time of the year) and for which the ice-water interface plays a crucial role in the lifecycle (75).

**Figure 6.**
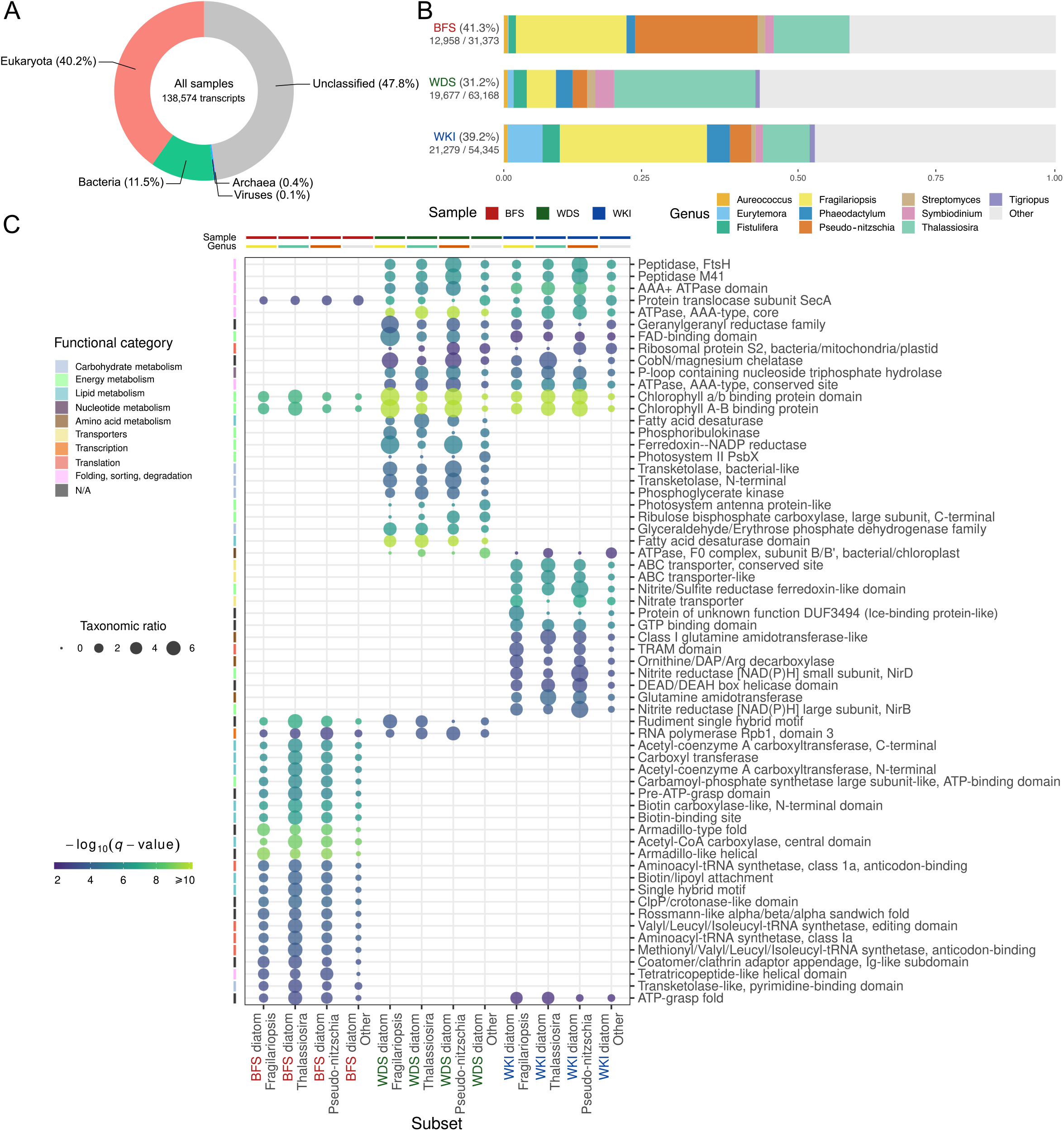
Diatom-rich communities metatranscriptome taxonomic classification and diatom-assigned transcript subsets InterPro enrichment results. (**A**) Domain-level taxonomic profile of the global metatranscriptome. 4,734 transcript sequences assigned to ‘cellular organisms’ or the root node of the taxonomy are not represented. (**B**) Genus-level taxonomic classification summary of transcripts expressed in each of the three sampling sites. For each sample, the fraction of expressed transcripts assigned to the top 10 represented genera (the genera to which the most transcripts were assigned to overall) is shown. Transcripts assigned to other less represented genera are aggregated as ‘Other’ (light grey fraction), and transcripts not assigned to any genus are not displayed. The numerical values complementing the sample identifiers indicate the ratio of transcripts classified at the genus-level over the total amount of expressed transcripts. (**C**) Heat map showing the 25 most enriched InterPro domains per subset of diatom-assigned transcripts expressed in each sampling site, compared with the global metatranscriptome (maximum enrichment *q*-value 0.01). Transcript subsets are represented along the x-axis and enriched InterPro domains along the y-axis. Transcripts associated to a subset and an enriched InterPro domain are binned by assigned genus (three most represented diatom genera and ‘Other’). For each combination of subset, enriched InterPro domain, and assigned genus, the circle size is proportional to the ‘taxonomic ratio’, a ratio of the frequency of the subset’s transcripts associated to the InterPro domain and assigned to the genus over the observed frequency for all the transcripts expressed in the sampling site. Enriched InterPro domains were assigned to broad functional categories, indicated by row annotations, and equivalent enrichment results were filtered to reduce redundancy. The significance of the enrichment is depicted as a color gradient, and each column is annotated with sample and genus classification information.

Functional enrichment analysis of diatom-assigned transcripts expressed in each sampling site sheds light on the functional variations between the diatom communities and reflects their adaptation to their respective environment (Figure 6C; Supplementary table S6-S8). Briefly, the two pelagic communities (BFS and WDS) were enriched for protein domains linked to carbohydrate and energy metabolic pathways, absent from the ice sheet community (WKI), potentially reflecting a reduced carbohydrate assimilation. InterPro entries linked to lipid metabolism were enriched in BFS (e.g. Acetyl-CoA carboxylase), potentially corresponding to nutrient stress in this community. Protein domains linked to translation and the basic protein processing machinery (e.g. peptidases, elongation factors, ribosomal proteins) or to membrane fluidity (e.g. fatty acid desaturase) were predominantly enriched in the two coldest communities, WDS and WKI, denoting a molecular footprint of their adaptation for survival at cold temperature. Finally, ice-binding proteins were enriched in WKI, with transcripts mainly assigned to *Fragilariopsis*, further alluding to the adaptation of this psychrophilic diatom to this stressful environment. Overall, these observations corroborate results obtained in a previous functional comparison study of these samples (14).

## CONCLUSIONS

We have presented TRAPID 2.0, a novel tool for the annotation and exploration of *de novo* assembled (meta)transcriptomes. Entirely web-based, it is well-suited for non-expert scientists as it removes the requirements for bioinformatics expertise or computational resources usually inherent to high-throughput –omics data processing. The high-quality reference databases it leverages enable the characterization of transcript sequences from a broad taxonomic range, while the components of its exploratory phase allow several downstream comparative and functional analyses. Detailed information and machine-readable export files are available for every processing or analysis step, ensuring reproducibility and facilitating subsequent analyses for advanced users.

Dissecting microbial eukaryote transcriptomes from the MMETSP with TRAPID 2.0 provided a glimpse into their diversity and complexity. Although they are bulk transcriptomes of single cultured species, the processed samples exhibited a wide array of taxonomic classification profiles, sequence counts, and gene space completeness scores. Their inspection additionally illustrated various features of TRAPID 2.0, showcasing the adaptability of its approach when confronted to the multifaceted nature of the analysis of transcriptomes. In practice, we have successfully used TRAPID 2.0 to generate complementary quality metrics, perform taxonomic, functional, and comparative analyses, and observe the impact of using an appropriate genetic code during ORF prediction.

Comparing TRAPID 2.0 to other similar transcriptome analysis software underscored its singularity and strengths, such as the inclusion of a built-in taxonomic classification module or its web-based analytical toolkit. Although TRAPID 2.0 does not perform transcript expression quantification, we have shown it is possible to exploit this information by defining and analyzing transcript subsets based on observed expression patterns. We have demonstrated TRAPID 2.0’s potential by employing it to annotate and analyze a metatranscriptome of three phytoplankton communities from the western Antarctica Peninsula, uncovering taxonomic and functional variations across distinct sampling sites.

In conclusion, TRAPID 2.0 provides researchers with a user-friendly and versatile tool to efficiently process *de novo* (meta)transcriptomes, and constitutes a valuable contribution to the ever-expanding transcriptome analysis software ecosystem.

## AVAILABILITY

TRAPID 2.0 can be reached at http://bioinformatics.psb.ugent.be/trapid. General documentation, tutorials, and example data sets are available on the TRAPID 2.0 website. The source code can be accessed online at https://github.com/VIB-PSB/trapid.

## SUPPLEMENTARY DATA

Supplementary Data are available at NAR online.

## ACKNOWLEDGEMENTS

We thank Luis Javier Galindo for his comments on the manuscript, and the early users of TRAPID 2.0 – Aurélien Carlier, the CNB research group, and the SINGEK network – for their avid testing of the platform, feedback, and suggestions.

