## Supplementary Data for "TRAPID 2.0: a web application for taxonomic and functional analysis of *de novo* transcriptomes"

#### SUPPLEMENTARY NOTES

##### Supplementary Note S1. Taxonomic classification evaluation

In line with other evaluation studies, taxonomic classification benchmark experiments were conducted performance was assessed using mock community data sets consisting of transcript sequences randomly sampled from a selection of phylogenetic clades as input, and excluding the input sequences from the reference indices.

After downloading all the RNA sequences from RefSeq (release 84, September 2017), four evaluation data sets were created. Each evaluation data set consist of a collection of 10,000 transcript sequences from mock communities, where sequences encompassed in selected phylogenetic clades are randomly sampled according to predefined proportions. Similarly to other sequence databases, RefSeq suffers from sampling bias in the phylogenetic distribution of available sequences: two types of evaluation data sets were therefore created to take that into consideration. The first one, named 'Minimal', contains sequences sampled from a small number of well-represented phylogenetic clades, for which many RNA sequences are available within RefSeq, and favors eukaryotic clades (Supplementary Table S9). The second one, named 'Order-100', aims to be less biased and more complex in its composition, and thus contain sequences sampled from all the orders for which 100 RNA sequences or more are available (244 orders in total), in equal proportions. Two replicates of each type of data set were generated and used to perform the benchmark experiments. All taxonomic classification benchmark experiments were performed using older versions of the NCBI taxonomy and non-redundant protein database than those currently used within TRAPID 2.0, that were both downloaded on the 18$^{th}$ of September 2017.

A total of five classification procedures were evaluated. Kaiju was run in MEM mode (using the default minimum fragment length of 11), as well as in Greedy mode (using the default minimum score s of 65), allowing up to five ('Greedy-5') amino acid substitutions/mismatches during the search, and always filtering low-complexity sequences. The same settings were replicated when Kaiju's index was split. To evaluate Kaiju's classification performance, we also executed a canonical protein sequence similarity search with DIAMOND, as a baseline method. DIAMOND was run in 'more sensitive' mode, using a maximum E-value cutoff of 1e-5, and the taxonomic classification of each aligned input sequence was considered to be the classification of the best hit. To avoid biasing evaluation results with self-hits, proteins corresponding to sequences present in the input data set were excluded from the Kaiju and DIAMOND indices while performing the taxonomic classification benchmarks, based on simple identifier mapping. Input transcript sequences corresponding to non-coding RNAs were ignored for the evaluation.

Taxonomic classification performance was evaluated at two taxonomic ranks: phylum and genus. For each considered taxonomic rank, sensitivity and precision were calculated as outlined in (1). Sensitivity and precision values determined for each evaluation setting were subsequently averaged across data set replicates to produce single values per classification procedure and data set type (Supplementary figure S1).

Our benchmarks confirmed the use of Kaiju for the classification of protein-coding transcripts: measured sensitivity and precision were in the range of the values reported in the evaluation reported in the original publication (figure 2 of (1), ‘454 single-end 350 nt’). Similar sensitivity and precision values were observed across the two types of input data sets (80.9-85.3% and 92.1-96.5% at phylum-level, respectively), although the classification of the ‘Order-100’ data set resulted in lower sensitivity. This difference was expected, given the paucity of closely related homologs in the reference data. For the tested benchmark data sets, DIAMOND performed better than Kaiju at the phylum level, yielding both higher sensitivity and precision, but not at the genus level where it achieved marginally better sensitivity at the cost of a lower precision than Kaiju, when considering all running modes. This observed lower precision may be due to the fact that only the top hit is taken into account to assign taxonomy. Another notable difference between the two programs is their execution time: Kaiju was reported to be 10 times faster than DIAMOND in the evaluation performed in (2). These results confirm Kaiju’s ability to successfully classify assembled transcript sequences and show that the running time required to run DIAMOND against a large reference database prohibits its use in TRAPID 2.0, where large sets of input sequences need to be processed. Comparing Kaiju running modes mirrored some of the results reported in the original publication: allowing 5 mismatches (‘Greedy-5’) resulted in better performance compared to the MEM mode at phylum-level, although in our setting the sensitivity gain was offset by a precision loss at genus-level. Finally, benchmark experiments confirmed that splitting Kaiju’s index into smaller chunks to lower its memory footprint resulted in nearly identical MEM mode performance, both at phylum and genus level. Therefore, Kaiju MEM with a split index was ultimately retained for the taxonomic classification step of TRAPID 2.0, since splitting the index did not result in a loss of performance and the MEM mode may give the best precision at genus-level, while it runs for a fraction of the time needed to perform an equivalent DIAMOND search.

**Supplementary Note S2. Functional annotation evaluation**

Functional annotation benchmark experiments were conducted using data from five model organisms: *Arabidopsis thaliana*, *Drosophila melanogaster*, *Escherichia coli* (strain K12)*, Homo sapiens*, and

*Saccharomyces cerevisiae* (strain ATCC 204508/S288c). Each organism's proteome GO annotation was retrieved from the GOA database (3) (unfiltered proteome annotation files, downloaded on 2018-11-15). The cDNA sequences corresponding to the proteins present in the annotation files were used as input for the benchmark experiments. To generate the input data sets, cDNA sequences were downloaded from Ensembl (4) and Ensembl genomes (5) (downloaded on 2018-11-15), and subsequently filtered using UniProt's ID mapping and Ensembl REST API cross-reference functionality to retain sequences matching with proteins from the GO annotation files exclusively. Each input data set was processed using TRAPID 2.0, selecting either PLAZA 4.5 dicots or eggNOG 4.5 as reference database. Taxonomic classification and non-coding RNA identification were disabled and all remaining initial processing parameters set to their default values. For PLAZA 4.5 dicots, the impact of the GF transfer cutoff, the minimum frequency among the members of a GF required for a functional annotation label to be assigned to a transcript set to 0.5 in TRAPID 2.0, was assessed. In total, 10 GF transfer cutoff were tested, ranging from 0 (i.e. the functional annotation labels associated to any member of the GF are transferred to the transcript) to 1 (i.e. only functional annotation labels that are associated to all the members of a GF are transferred to the transcript).

For each cDNA sequence, functional annotation performance was evaluated by comparing the set of predicted GO terms to the 'gold standard' GO terms, here defined as the GO terms associated to the corresponding protein with experimental, IC (inferred by curator), or TAS (traceable author statement) GO evidence codes. Sensitivity, precision, and $F_1$ score were computed as described in (6), for each GO aspect. To produce single evaluation values per organism, GO aspect, and experimental setting, the metrics computed for individual cDNA sequences were averaged across all the sequences of an organism (ignoring those for which evaluation metrics were impossible to get). In addition to the functional annotation performance metrics, the annotation coverage, corresponding to the fraction of sequences for which GO terms were predicted, was also computed.

To prevent biasing the functional annotation evaluation metrics with frequent and broad GO terms not carrying a lot of information, 86 GO terms were filtered out and ignored in the benchmark experiments. The set of filtered GO terms consists of the three root GO terms ('cellular_component', 'molecular_function', 'biological_process'), and 83 GO terms that were associated to at least 20% of the proteins of any of the considered organisms in the 'true' GO annotations.

#### SUPPLEMENTARY FIGURES

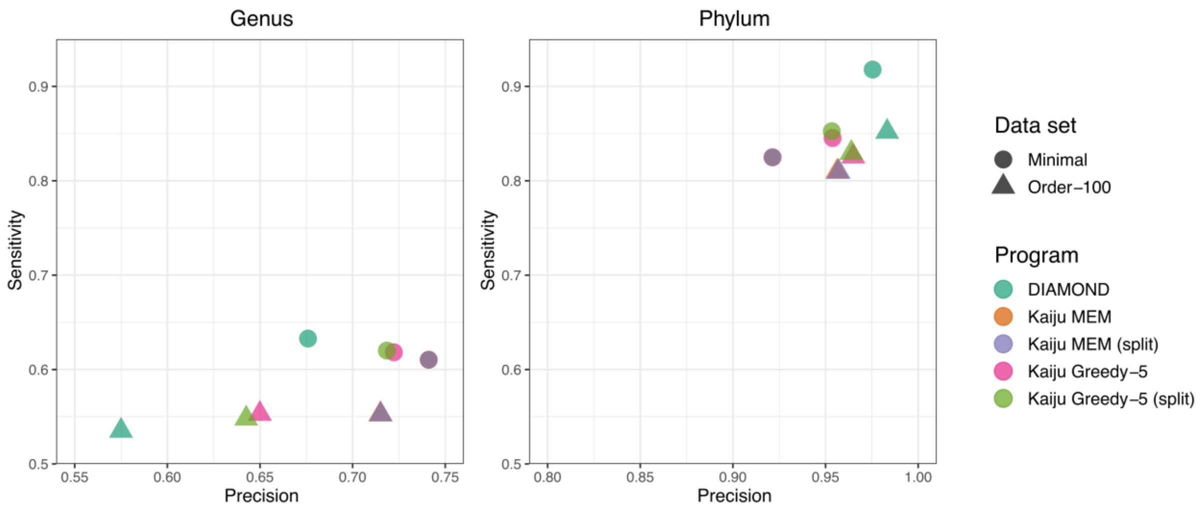


**Supplementary figure S1.** Taxonomic classification and genus- and phylum-level sensitivity and precision. For all evaluated programs, sensitivity and precision values were averaged over replicates of each type of evaluation data set.

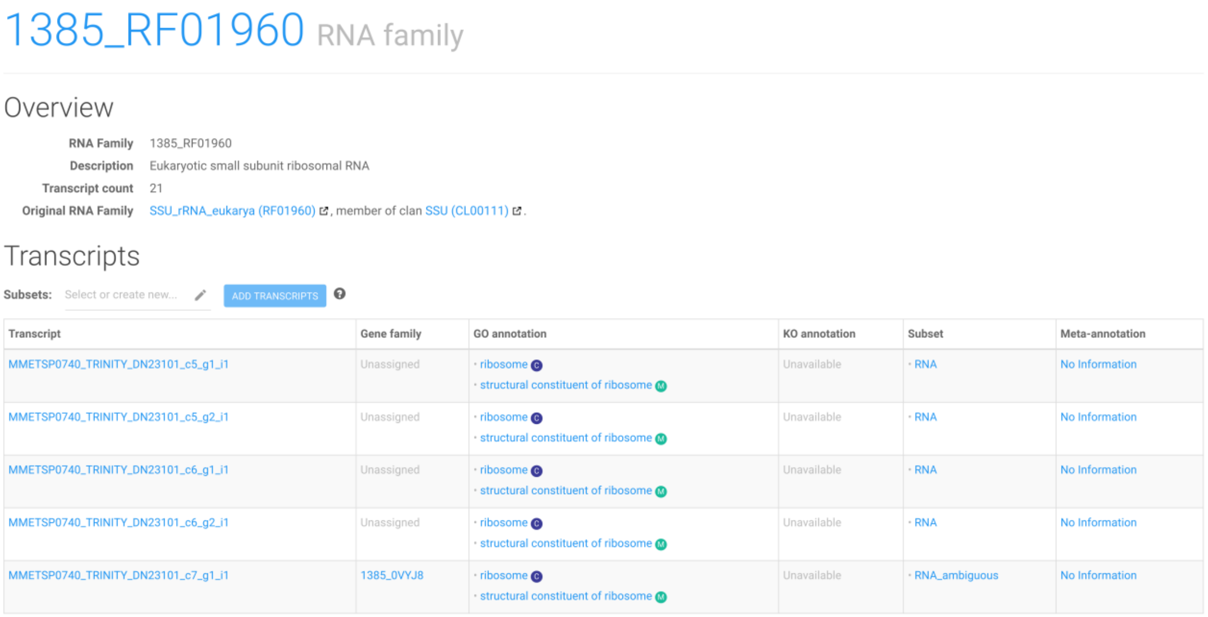

### 1385_RF01960 RNA family

#### Overview

**RNA Family** 1385_RF01960
**Description** Eukaryotic small subunit ribosomal RNA
**Transcript count** 21
**Original RNA Family** SSU_rRNA_eukarya (RF01960), member of clan SSU (CL00111).

#### Transcripts

**Subsets:** Select or create new... ADD TRANSCRIPTS

| Transcript | Gene family | GO annotation | KO annotation | Subset | Meta-annotation |
|---|---|---|---|---|---|
| MMETSP0740_TRINITY_DN23101_c5_g1_i1 | Unassigned | · ribosome C<br>· structural constituent of ribosome M | Unavailable | · RNA | No Information |
| MMETSP0740_TRINITY_DN23101_c5_g2_i1 | Unassigned | · ribosome C<br>· structural constituent of ribosome M | Unavailable | · RNA | No Information |
| MMETSP0740_TRINITY_DN23101_c6_g1_i1 | Unassigned | · ribosome C<br>· structural constituent of ribosome M | Unavailable | · RNA | No Information |
| MMETSP0740_TRINITY_DN23101_c6_g2_i1 | Unassigned | · ribosome C<br>· structural constituent of ribosome M | Unavailable | · RNA | No Information |
| MMETSP0740_TRINITY_DN23101_c7_g1_i1 | 1385_0VYJ8 | · ribosome C<br>· structural constituent of ribosome M | Unavailable | · RNA_ambiguous | No Information |

**Supplementary figure S2. RNA family page screenshot.** Twenty-one transcripts, listed in the table, were assigned to the ‘RF01960’ RNA family (‘SSU_rRNA_eukarya’). Links to Rfam pages for this family and its encompassing clan are available, and further investigation of individual transcripts is possible. The form on top of the table enables the addition of transcripts to an existing or new subset. GO terms associated to the RNA family were transferred to the transcript sequences (‘GO annotation’ column). Only five transcripts of the table are shown to improve legibility. These results were generated using MMETSP0740 (*Thalassiosira minuscula* CCMP1093) with eggNOG 4.5 as a reference database and default initial processing parameters.

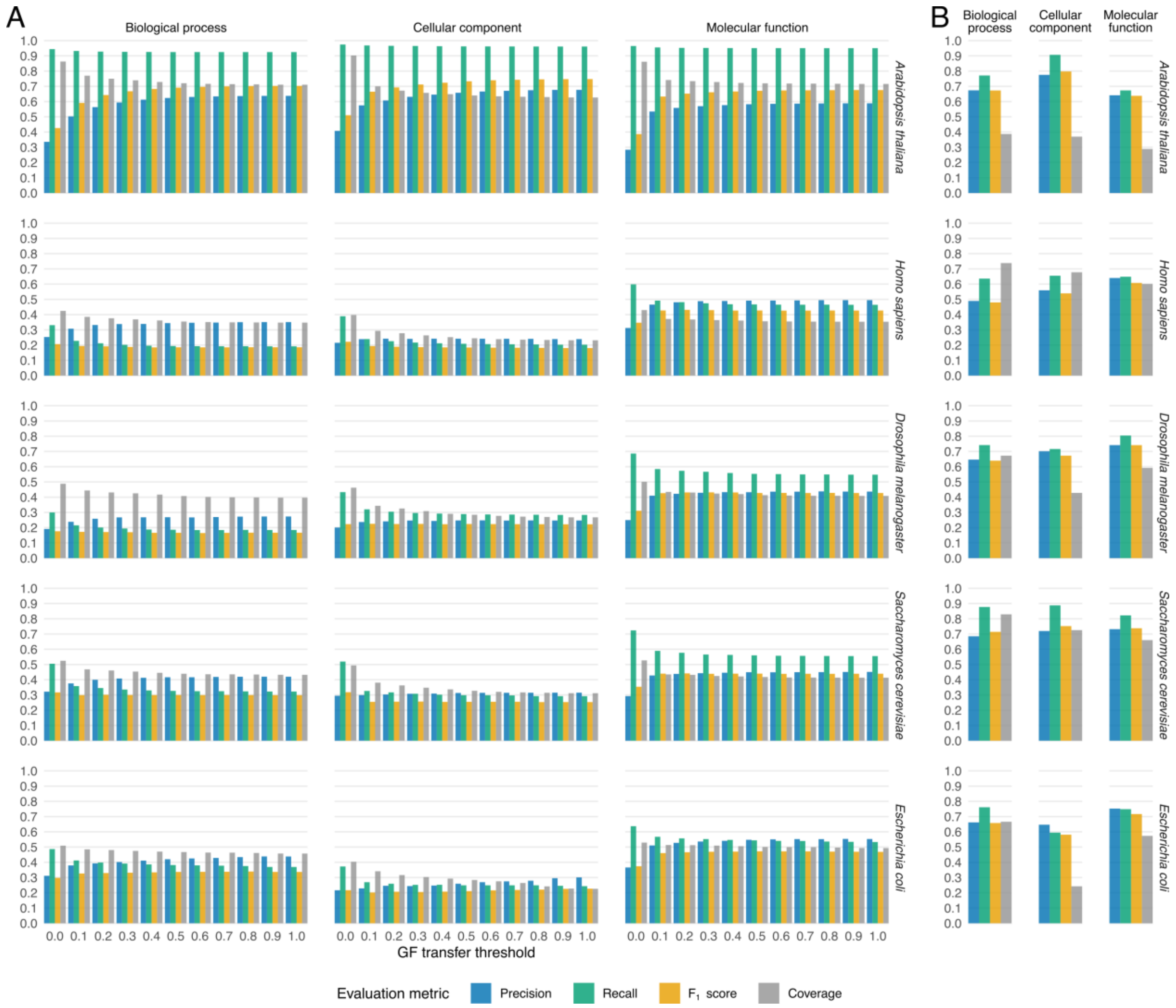


**Supplementary figure S3.** TRAPID 2.0 GO annotation performance evaluation. GO annotation evaluation metrics yielded when selecting PLAZA 4.5 dicots as a reference database (**A**), testing varying gene family transfer thresholds, or eggNOG 4.5 (**B**). Each row corresponds to one of the five considered model organisms, and each column to a GO aspect.

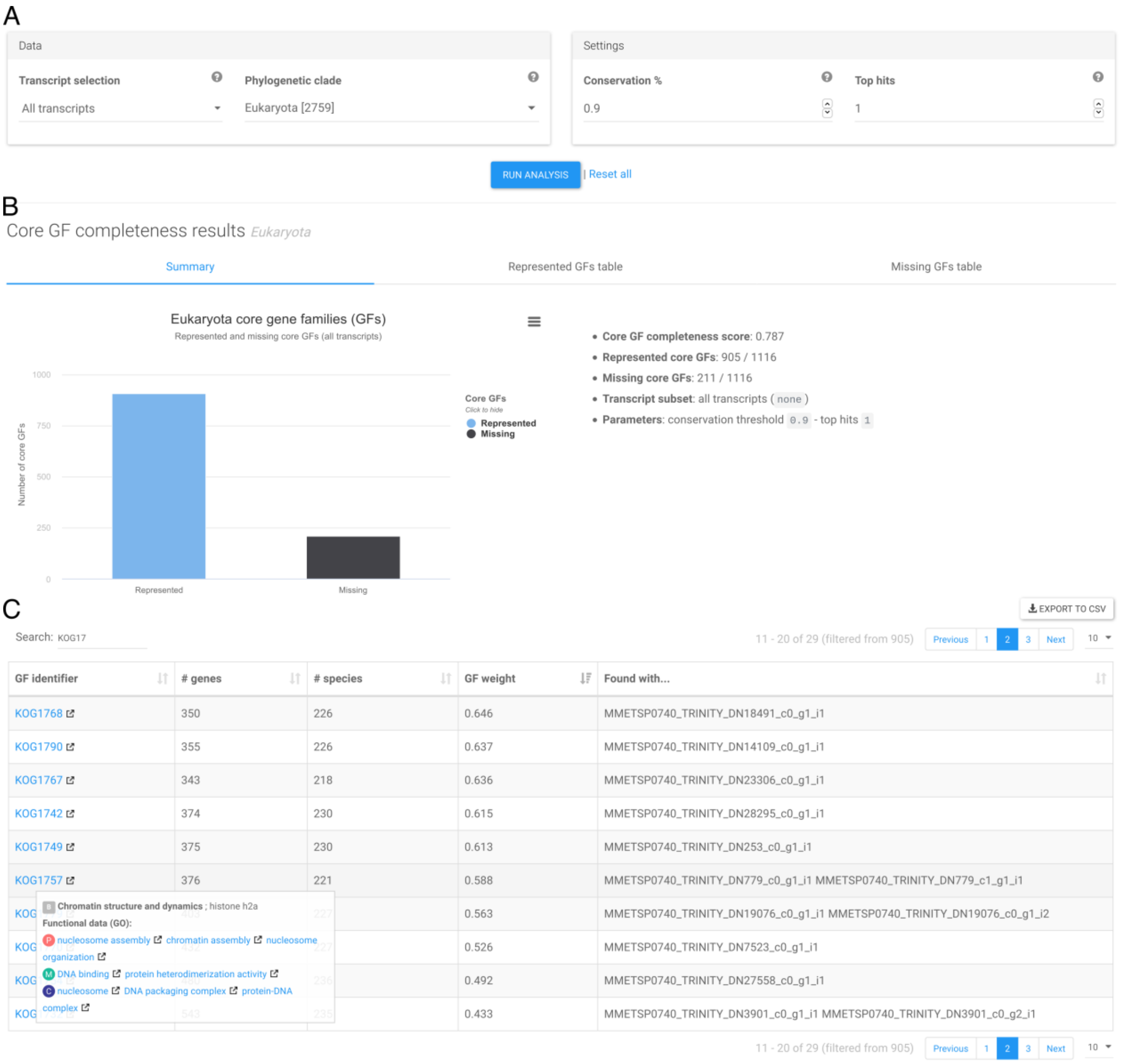


**Supplementary figure S4.** Core gene family completeness analysis module screenshot. (**A**) Job submission panel. Any phylogenetic clade represented in the selected reference database can be used for the analysis. The species conservation threshold, used to define core gene families, is also adjustable to fit variable stringency requirements. (**B**) Analysis results panel. Results are summarized as a bar chart depicting the number of represented and missing core gene families, the completeness score and additional analysis metrics. (**C**) Represented core gene families table. Represented or missing core gene families and their associated functional data can be further investigated using the dedicated tables. These results were generated using MMETSP0740 (*Thalassiosira minuscula* CCMP1093) with eggNOG 4.5 as a reference database.

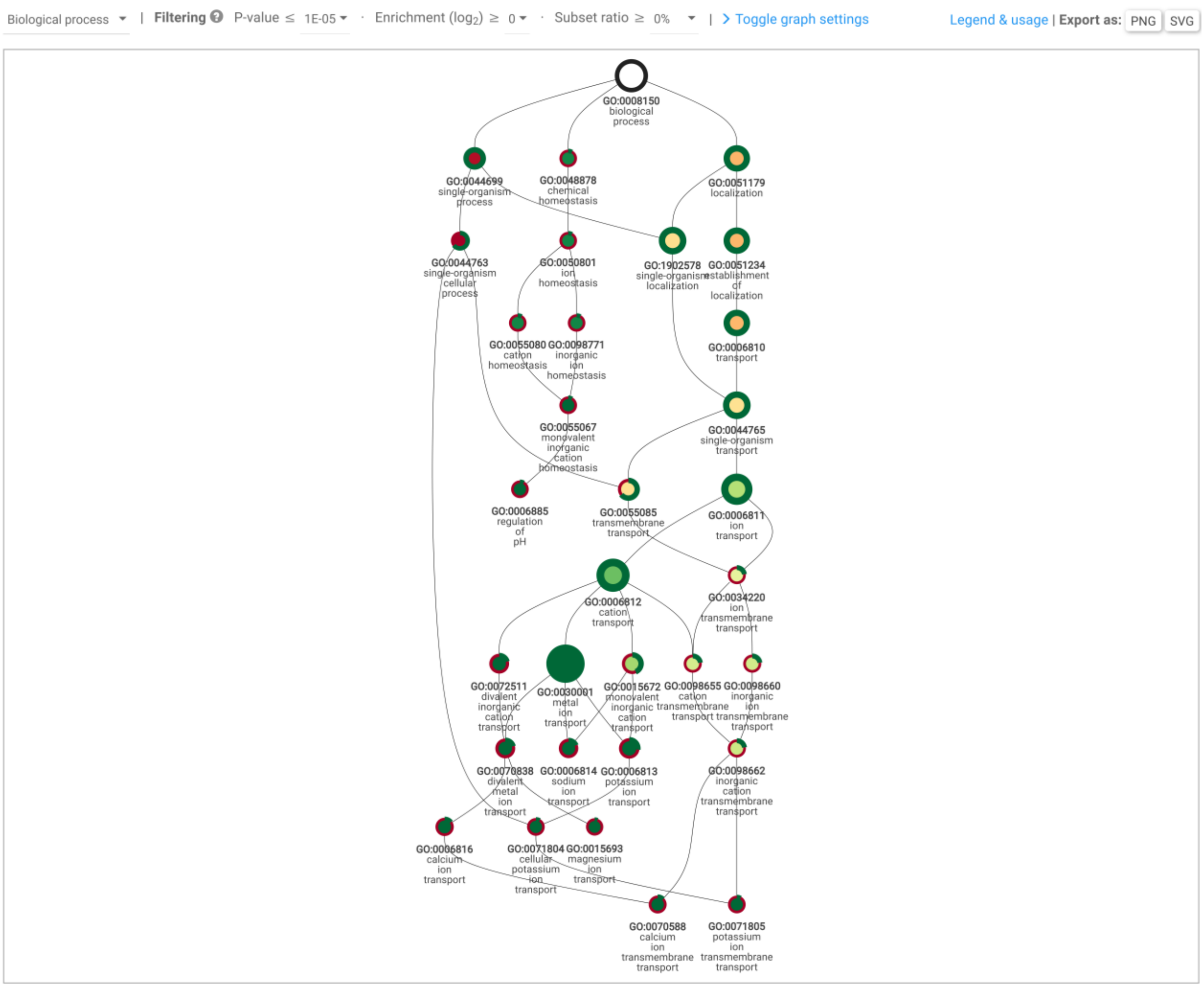


**Supplementary figure S5.** Interactive subgraph of enriched GO terms for a subset of 48 *Ostreococcus mediterraneus* metal ion transport transcripts. The nodes represent enriched GO terms and are arranged hierarchically, with ancestral terms (more general) displayed on top and descendant terms (more specialized) at the bottom. The node size is proportional to the GO term's enrichment *q*-value, with a larger size indicating a lower *q*-value (i.e., a more significant enrichment). The node color corresponds to the GO term's enrichment fold, ranging from red (lowest) to green (highest). The node outline depends on the number of transcripts annotated with the GO term in the subset: the green part represents the fraction of transcripts in which it is present and the red part the fraction in which it is absent. The control panel displayed on top of the graph enables the user to choose the GO aspect to visualize (here biological process), to filter nodes based on minimum or maximum values of enrichment metrics, and to adjust the appearance of the graph. Clicking any GO identifier on the graph opens the dedicated GO term page. Hovering over a node displays a tooltip detailing the corresponding enrichment metrics, and right-clicking it shows a menu with node selection and hiding options. The maximum enrichment *q*-value threshold is 1e-5. These results were generated using MMETSP0936 (*Ostreococcus mediterraneus* clade-D-RCC2573) with pico-PLAZA 3 as a reference database and default initial processing parameters.

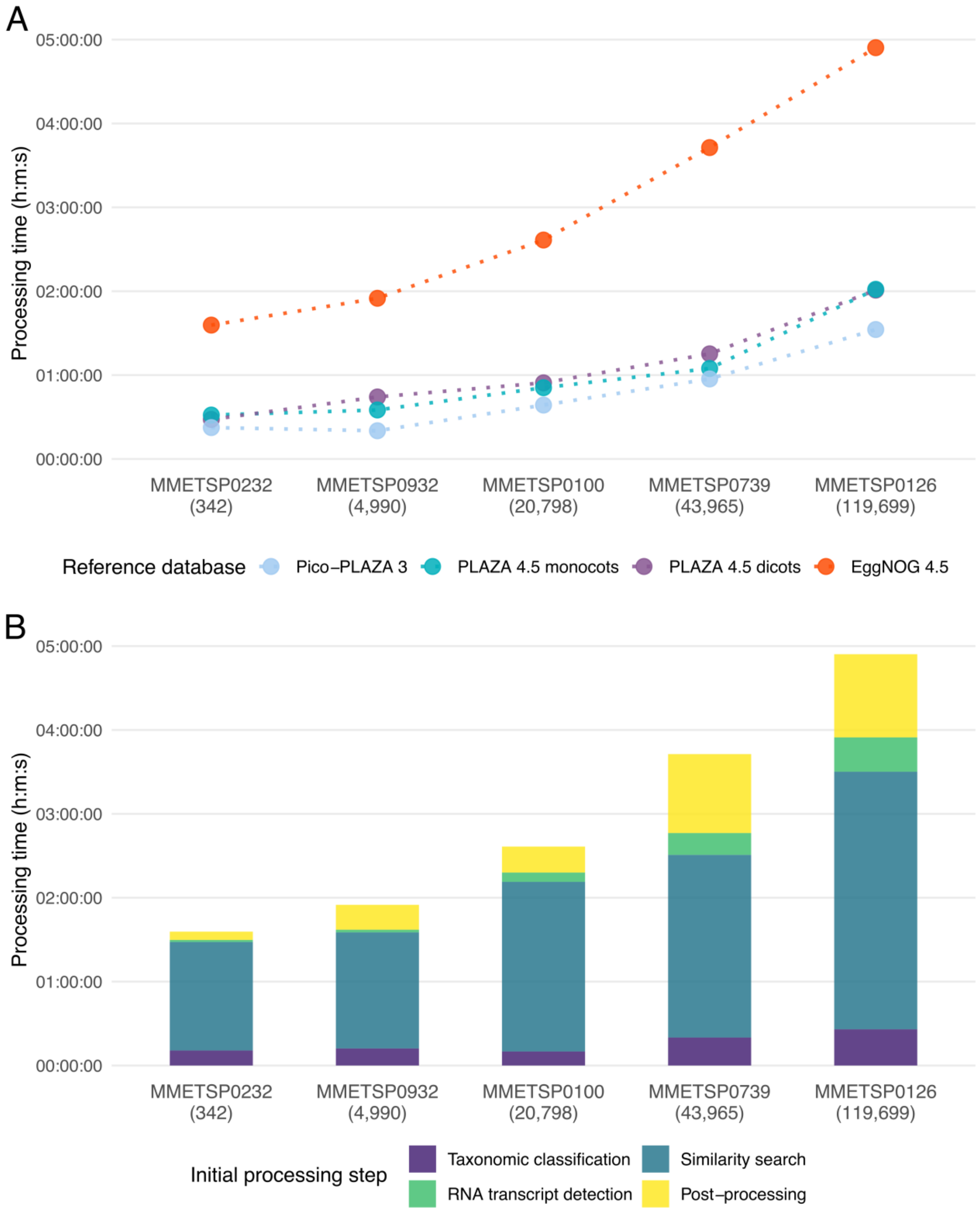


**Supplementary figure S6.** TRAPID 2.0 initial processing execution time for five MMETSP transcriptomes. **(A)** Total initial processing execution time for all tested transcriptomes and available reference databases. **(B)** Execution time per initial processing step, for all tested transcriptomes and using eggNOG 4.5 as reference database. The transcript count of each transcriptome is indicated between brackets. All initial processing parameters were set to their default values. The reported execution times are average values obtained over four runs.

#### SUPPLEMENTARY TABLES

**Supplementary Table S1.** Sample name, transcript count, and taxonomy of processed MMETSP transcriptomes. Sample taxonomy information was retrieved from the MMETSP metadata.

| Sample name | #transcripts | Phylum | Genus | Species | Strain |
|---|---|---|---|---|---|
| **MMETSP0018** | 19,640 | Ciliophora | Uronema | sp. | Bbcil |
| **MMETSP0098** | 45,351 | Unknown | Unknown | Unknown | NY0313808BC1 |
| **MMETSP0099** | 17,555 | Unknown | Unknown | Unknown | NY0313808BC1 |
| **MMETSP0100** | 20,798 | Unknown | Unknown | Unknown | NY0313808BC1 |
| **MMETSP0123** | 38,054 | Ciliophora | Favella | ehrenbergii | Fehren 1 |
| **MMETSP0125** | 41,818 | Ciliophora | Aristerostoma | sp. | ATCC 50986 |
| **MMETSP0126** | 119,699 | Ciliophora | Strombidinopsis | acuminatum | SPMC142 |
| **MMETSP0127** | 83,965 | Ciliophora | Platyophrya | macrostoma | WH |
| **MMETSP0208** | 42,764 | Ciliophora | Strombidium | inclinatum | S3 |
| **MMETSP0211** | 44,971 | Ciliophora | Pseudokeronopsis | sp. | OXSARD2 |
| **MMETSP0216** | 65,706 | Ciliophora | Protocruzia | adherens | Boccale |
| **MMETSP0232** | 342 | Ascomycota | Debaryomyces | hansenii | J26 |
| **MMETSP0233** | 9,596 | Ascomycota | Debaryomyces | hansenii | J26 |
| **MMETSP0288** | 45,732 | Apicomplexa | Alveolata | sp. | CCMP3155 |
| **MMETSP0434** | 48,003 | Ciliophora | Favella | taraikaensis | Fe Narragansett Bay |
| **MMETSP0436** | 33,913 | Ciliophora | Favella | taraikaensis | Fe Narragansett Bay |
| **MMETSP0449** | 19,501 | Ciliophora | Strombidium | rassoulzadegani | ras09 |
| **MMETSP0463** | 43,507 | Ciliophora | Strombidinopsis | sp | SopsisLIS2011 |
| **MMETSP0472** | 135,916 | Ciliophora | Tiarina | fusus | LIS |
| **MMETSP0737** | 47,167 | Bacillariophyta | Thalassiosira | miniscula | CCMP1093 |
| **MMETSP0738** | 34,790 | Bacillariophyta | Thalassiosira | miniscula | CCMP1093 |
| **MMETSP0739** | 43,965 | Bacillariophyta | Thalassiosira | miniscula | CCMP1093 |
| **MMETSP0740** | 36,324 | Bacillariophyta | Thalassiosira | miniscula | CCMP1093 |
| **MMETSP0929** | 9,005 | Chlorophyta | Ostreococcus | mediterraneus | clade-D-RCC2572 |
| **MMETSP0930** | 9,482 | Chlorophyta | Ostreococcus | mediterraneus | clade-D-RCC1621 |
| **MMETSP0932** | 4,990 | Chlorophyta | Ostreococcus | mediterraneus | clade-D-RCC2596 |
| **MMETSP0936** | 9,258 | Chlorophyta | Ostreococcus | mediterraneus | clade-D-RCC2573 |
| **MMETSP0937** | 9,289 | Chlorophyta | Ostreococcus | mediterraneus | clade-D-RCC2593 |
| **MMETSP0938** | 9,193 | Chlorophyta | Ostreococcus | mediterraneus | clade-D-RCC1107 |
| **MMETSP1018** | 17,679 | Ciliophora | Anophryoides | haemophila | AH6 |
| **MMETSP1019** | 21,468 | Ciliophora | Anophryoides | haemophila | AH6 |
| **MMETSP1396** | 49,169 | Ciliophora | Pseudokeronopsis | sp. | Brazil |
| **MMETSP1451** | 54,204 | Alveolata | Vitrella | brassicaformis | CCMP3346 |
| **MMETSP1456** | 12,219 | Unknown | Unidentified | sp. | RCC1871 |

**Supplementary Table S2.** Overview of TRAPID 2.0 reference databases. The gene family count only includes homology-based families for PLAZA databases, and only orthologous groups at the root level for eggNOG 4.5.

| Reference database | #Species | #Genes | #Gene families | Taxonomic focus | Functional annotation | Gene family construction |
|---|---|---|---|---|---|---|
| PLAZA 4.5 dicots | 55 | 3,065,012 | 208,456 | Dicot plants | GO, InterPro | Tribe-MCL, integrative orthologs |
| PLAZA 4.5 monocots | 39 | 1,563,555 | 213,318 | Monocot plants | GO, InterPro | Tribe-MCL, integrative orthologs |
| pico-PLAZA 3 | 39 | 705,020 | 127,718 | Microbial photosynthetic eukaryotes | GO, InterPro | Tribe-MCL, integrative orthologs |
| eggNOG 4.5 | 2,031 | 9,646,196 | 190,803 | Archaea, Bacteria, Eukaryotes | GO, KO | eggNOG |

**Supplementary Table S3.** Read filtering and mapping information for three Antarctic Peninsula phytoplankton community metatranscriptomes from the Bransfield Strait (BFS), Wilkins sea ice (WKI) and Weddell Sea (WDS) samples. Raw reads from the three samples were filtered and trimmed, and used for metatranscriptome assembly and expression quantification. The values reported in the '# Suppl. Alignments' and the '#Unmapped in unused' columns correspond to the number of mapped reads having supplementary alignments, and the number of unmapped reads that are in data that is unused in the assembly, respectively.

| Source | SRA accession | #Raw reads | #Quality-filtered reads (% raw reads) | #Mapped (% retained reads) | #Suppl. alignments (% retained reads) | #Unmapped (% retained reads) | #Unmapped in unused (% retained reads) |
|---|---|---|---|---|---|---|---|
| BFS | SRR1606327 | 114,323 | 108,291 (0.95) | 88,426 (81.66) | 174 (0.16) | 19,865 (18.34) | 18,100 (16.71) |
| WDS | SRR1606329 | 253,631 | 239,196 (0.94) | 185,275 (77.46) | 375 (0.16) | 53,921 (22.54) | 48,809 (20.41) |
| WKI | SRR1606328 | 191,057 | 179,040 (0.94) | 137,764 (76.95) | 183 (0.10) | 41,276 (23.05) | 37,333 (20.85) |
| Total | - | 559,011 | 526,527 (0.94) | 411,465 (78.15) | 732 (0.14) | 115,062 (21.85) | 104,242 (19.80) |

**Supplementary Table S4.** Overview of all TRAPID 2.0 processing steps and their purpose. Processing steps are organized by processing phase, initial processing or exploratory, and chronological execution order when possible. Indentation indicates hierarchical relationships between processing steps.

| Processing phase | Processing step | Purpose |
|---|---|---|
| **Initial processing phase** | | Global transcriptome characterization |
| | Taxonomic classification | Potential contaminant detection; characterization of communities |
| | RNA transcripts detection | Non-coding RNA identification and annotation |
| | Sequence similarity search | Identification of homologous sequences within available reference databases |
| | Sequence similarity search post-processing | Characterization of protein-coding transcripts |
| | ⌞ Gene Family assignment | Clustering of transcripts with their homologs |
| | ⌞ Functional annotation transfer | Functional annotation of transcript sequences |
| | ⌞ Meta-annotation inference | Estimation of the fragmentation state of transcripts |
| | ⌞ Frame detection and ORF finding | Putative frameshift detection; coding sequence detection and translation (supporting non-canonical genetic codes) |
| **Exploratory phase** | | Refined comparative and functional analyses; examination of selected data types |
| | Experiment statistics | Exploration of annotation metrics and sequence length distribution |
| | Global protein alignment and phylogenetic tree construction | Analysis of transcripts and their homologs in a gene-family and evolutionary context; identification of orthology and paralogy relationships |
| | Core GF completeness analysis | Gene space completeness assessment and analysis at multiple phylogenetic levels |
| | Subset functional enrichment analysis | Identification and analysis of functional biases; discovery and exploration of significantly over-represented biological functions |
| | Subset functional annotation comparison | Exploration of the biological function of transcript subsets; comparison of functional biases |

**Supplementary Table S5.** TRAPID 2.0 experiment statistics summary for the global Antarctic Peninsula phytoplankton community metatranscriptome.

| Metric | Value |
|---|---|
| **Transcript information** | |
| #Transcripts | 143,308 |
| Average sequence length | 308 |
| Average ORF length (% total) | 245.9 |
| #Partial (% total) | 44,248 (30.9%) |
| **Taxonomic classification information** | |
| #Classified (% total) | 77,105 (53.8%) |
| #Unclassified (% total) | 66,203 (46.2%) |
| **Similarity search information (top 5)** | |
| Fragilariopsis cylindrus (% total) | 16,908 (26.57%) |
| Thalassiosira pseudonana (% total) | 13,796 (21.68%) |
| Phaeodactylum tricornutum (% total) | 8,919 (14.01%) |
| Drosophila melanogaster (% total) | 2,651 (4.17%) |
| Emiliania huxleyi (% total) | 2,132 (3.35%) |
| Total hits (% total) | 63,641 (44.41%) |
| **Gene family information** | |
| #Gene families | 8,931 |
| #Transcripts in GF (% total) | 63,641 (44.4%) |
| Largest GF | HOM03P000062 (602 transcripts) |
| #Single copy | 3,409 |
| **RNA family information** | |
| #RNA families | 13 |
| #Transcripts in RF (% total) | 4,536 (3.2%) |
| Largest RF | RF02543 (1498 transcripts) |
| **Functional annotation information** | |
| #GO terms | 12,965 |
| #Transcripts with GO (% total) | 65,831 (45.9%) |
| #InterPro domains | 7,052 |
| #Transcripts with Protein Domain (% total) | 63,641 (44.4%) |

**Supplementary Table S6.** Enriched InterPro domains in diatom-assigned transcripts expressed in the Bransfield Strait (BFS) sample, compared with the whole Antarctic Peninsula phytoplankton community metatranscriptome. Enrichment fold values are expressed in $\log_2$ scale and the subset ratio indicates the fraction of transcripts annotated with the enriched functional annotation label within the subset. Maximum enrichment *q*-value threshold is 0.01.

| Identifier | *q*-value | Enrichment fold | Subset ratio | Description |
|---|---|---|---|---|
| IPR011989 | 6.568E-10 | 0.733 | 2.875 | Armadillo-like helical |
| IPR013537 | 1.595E-09 | 2.117 | 0.390 | Acetyl-CoA carboxylase, central domain |
| IPR016024 | 2.301E-09 | 0.610 | 3.742 | Armadillo-type fold |
| IPR023329 | 4.040E-08 | 0.633 | 3.005 | Chlorophyll a/b binding protein domain |
| IPR022796 | 5.224E-08 | 0.632 | 2.947 | Chlorophyll A-B binding protein |
| IPR011054 | 8.971E-08 | 1.372 | 0.693 | Rudiment single hybrid motif |
| IPR005481 | 3.978E-07 | 1.608 | 0.477 | Biotin carboxylase-like, N-terminal domain |
| IPR005482 | 3.978E-07 | 1.608 | 0.477 | Biotin carboxylase, C-terminal |
| IPR011764 | 3.978E-07 | 1.608 | 0.477 | Biotin carboxylation domain |
| IPR001882 | 4.484E-07 | 1.591 | 0.477 | Biotin-binding site |
| IPR000022 | 1.302E-06 | 1.717 | 0.390 | Carboxyl transferase |
| IPR011763 | 1.761E-06 | 1.695 | 0.390 | Acetyl-coenzyme A carboxyltransferase, C-terminal |
| IPR011762 | 2.376E-06 | 1.674 | 0.390 | Acetyl-coenzyme A carboxyltransferase, N-terminal |
| IPR005479 | 2.628E-06 | 1.235 | 0.665 | Carbamoyl-phosphate synthetase large subunit-like, ATP-binding domain |
| IPR016185 | 4.922E-06 | 1.103 | 0.780 | Pre-ATP-grasp domain |
| IPR029045 | 7.531E-05 | 1.021 | 0.737 | ClpP/crotonase-like domain |
| IPR000089 | 1.067E-04 | 1.113 | 0.607 | Biotin/lipoyl attachment |
| IPR009080 | 1.119E-04 | 1.072 | 0.650 | Aminoacyl-tRNA synthetase, class 1a, anticodon-binding |
| IPR011053 | 1.215E-04 | 1.104 | 0.607 | Single hybrid motif |
| IPR002300 | 1.565E-04 | 1.313 | 0.433 | Aminoacyl-tRNA synthetase, class Ia |
| IPR014729 | 1.805E-04 | 0.642 | 1.560 | Rossmann-like alpha/beta/alpha sandwich fold |
| IPR009008 | 1.842E-04 | 1.325 | 0.419 | Valyl/Leucyl/Isoleucyl-tRNA synthetase, editing domain |
| IPR013155 | 2.144E-04 | 1.310 | 0.419 | Methionyl/Valyl/Leucyl/Isoleucyl-tRNA synthetase, anticodon-binding |
| IPR013041 | 6.156E-04 | 1.443 | 0.318 | Coatomer/clathrin adaptor appendage, Ig-like subdomain |
| IPR011990 | 6.323E-04 | 0.482 | 2.355 | Tetratricopeptide-like helical domain |
| IPR005475 | 8.717E-04 | 0.998 | 0.592 | Transketolase-like, pyrimidine-binding domain |
| IPR011761 | 8.945E-04 | 0.812 | 0.852 | ATP-grasp fold |
| IPR001412 | 9.054E-04 | 0.944 | 0.650 | Aminoacyl-tRNA synthetase, class I, conserved site |
| IPR013815 | 1.208E-03 | 0.836 | 0.780 | ATP-grasp fold, subdomain 1 |
| IPR002553 | 1.269E-03 | 1.222 | 0.390 | Clathrin/coatomer adaptor, adaptin-like, N-terminal |
| IPR002885 | 1.281E-03 | 0.955 | 0.592 | Pentatricopeptide repeat |
| IPR000185 | 1.367E-03 | 1.321 | 0.332 | Protein translocase subunit SecA |
| IPR011115 | 1.367E-03 | 1.321 | 0.332 | SecA DEAD-like, N-terminal |
| IPR011116 | 1.367E-03 | 1.321 | 0.332 | SecA Wing/Scaffold |
| IPR011130 | 1.367E-03 | 1.321 | 0.332 | SecA preprotein, cross-linking domain |
| IPR014018 | 1.367E-03 | 1.321 | 0.332 | SecA motor DEAD |
| IPR020937 | 1.367E-03 | 1.321 | 0.332 | SecA conserved site |
| IPR001478 | 1.395E-03 | 0.860 | 0.708 | PDZ domain |
| IPR017932 | 1.425E-03 | 1.016 | 0.535 | Glutamine amidotransferase type 2 domain |
| IPR003440 | 1.510E-03 | 2.080 | 0.144 | Glycosyl transferase, family 48 |
| IPR026899 | 1.510E-03 | 2.080 | 0.144 | 1,3-beta-glucan synthase subunit FKS1-like, domain-1 |
| IPR007066 | 1.593E-03 | 1.076 | 0.462 | RNA polymerase Rpb1, domain 3 |
| IPR007081 | 1.593E-03 | 1.076 | 0.462 | RNA polymerase Rpb1, domain 5 |
| IPR007083 | 1.593E-03 | 1.076 | 0.462 | RNA polymerase Rpb1, domain 4 |
| IPR029061 | 2.251E-03 | 0.784 | 0.780 | Thiamin diphosphate-binding fold |
| IPR019734 | 2.282E-03 | 0.604 | 1.257 | Tetratricopeptide repeat |
| IPR009014 | 4.751E-03 | 0.928 | 0.520 | Transketolase, C-terminal/Pyruvate-ferredoxin oxidoreductase, domain II |
| IPR005467 | 6.044E-03 | 1.199 | 0.318 | Histidine kinase domain |
| IPR005113 | 6.123E-03 | 1.487 | 0.217 | uDENN domain |
| IPR001194 | 6.430E-03 | 1.421 | 0.231 | DENN domain |

| | | | | |
|---|---|---|---|---|
| IPR001296 | 6.468E-03 | 1.765 | 0.159 | Glycosyl transferase, family 1 |
| IPR000403 | 6.510E-03 | 1.132 | 0.347 | Phosphatidylinositol 3-/4-kinase, catalytic domain |
| IPR010788 | 7.233E-03 | 2.100 | 0.116 | VDE lipocalin domain |
| IPR001108 | 9.669E-03 | 2.421 | 0.087 | Peptidase A22A, presenilin |

**Supplementary Table S7.** Enriched InterPro domains in diatom-assigned transcripts expressed in the Weddell Sea (WDS) sample, compared with the whole Antarctic Peninsula phytoplankton community metatranscriptome. Enrichment fold values are expressed in $log_2$ scale and the subset ratio indicates the fraction of transcripts annotated with the enriched functional annotation label within the subset. Maximum enrichment *q*-value threshold is 0.01.

| **Identifier** | ***q*-value** | **Enrich-ment fold** | **Subset ratio** | **Description** |
|---|---|---|---|---|
| IPR023329 | 9.883E-60 | 1.278 | 4.699 | Chlorophyll a/b binding protein domain |
| IPR022796 | 9.554E-58 | 1.269 | 4.583 | Chlorophyll A-B binding protein |
| IPR003959 | 1.407E-11 | 0.836 | 2.151 | ATPase, AAA-type, core |
| IPR005804 | 1.041E-10 | 1.347 | 0.832 | Fatty acid desaturase domain |
| IPR002146 | 2.015E-09 | 2.118 | 0.320 | ATPase, F0 complex, subunit B/B', bacterial/chloroplast |
| IPR000185 | 2.438E-07 | 1.623 | 0.410 | Protein translocase subunit SecA |
| IPR011115 | 2.438E-07 | 1.623 | 0.410 | SecA DEAD-like, N-terminal |
| IPR011116 | 2.438E-07 | 1.623 | 0.410 | SecA Wing/Scaffold |
| IPR011130 | 2.438E-07 | 1.623 | 0.410 | SecA preprotein, cross-linking domain |
| IPR014018 | 2.438E-07 | 1.623 | 0.410 | SecA motor DEAD |
| IPR020937 | 2.438E-07 | 1.623 | 0.410 | SecA conserved site |
| IPR006424 | 4.081E-07 | 1.056 | 0.845 | Glyceraldehyde-3-phosphate dehydrogenase, type I |
| IPR020828 | 4.081E-07 | 1.056 | 0.845 | Glyceraldehyde 3-phosphate dehydrogenase, NAD(P) binding domain |
| IPR020829 | 4.081E-07 | 1.056 | 0.845 | Glyceraldehyde 3-phosphate dehydrogenase, catalytic domain |
| IPR020830 | 4.081E-07 | 1.056 | 0.845 | Glyceraldehyde 3-phosphate dehydrogenase, active site |
| IPR020831 | 4.081E-07 | 1.056 | 0.845 | Glyceraldehyde/Erythrose phosphate dehydrogenase family |
| IPR000642 | 1.002E-06 | 1.319 | 0.525 | Peptidase M41 |
| IPR000685 | 1.031E-06 | 1.865 | 0.282 | Ribulose bisphosphate carboxylase, large subunit, C-terminal |
| IPR017443 | 1.031E-06 | 1.865 | 0.282 | Ribulose bisphosphate carboxylase, large subunit, ferrodoxin-like N-terminal |
| IPR020878 | 1.031E-06 | 1.865 | 0.282 | Ribulose bisphosphate carboxylase, large chain, active site |
| IPR005936 | 1.378E-06 | 1.321 | 0.512 | Peptidase, FtsH |
| IPR000932 | 1.700E-06 | 1.258 | 0.551 | Photosystem antenna protein-like |
| IPR003593 | 6.505E-06 | 0.388 | 4.391 | AAA+ ATPase domain |
| IPR015701 | 6.701E-06 | 1.578 | 0.333 | Ferredoxin--NADP reductase |
| IPR002938 | 1.788E-05 | 1.247 | 0.474 | FAD-binding domain |
| IPR027417 | 1.851E-05 | 0.210 | 12.252 | P-loop containing nucleoside triphosphate hydrolase |
| IPR006082 | 1.932E-05 | 1.795 | 0.243 | Phosphoribulokinase |
| IPR012171 | 3.614E-05 | 1.466 | 0.333 | Fatty acid desaturase |
| IPR005478 | 7.070E-05 | 1.313 | 0.384 | Transketolase, bacterial-like |
| IPR009518 | 7.504E-05 | 2.832 | 0.090 | Photosystem II PsbX |
| IPR005474 | 8.104E-05 | 1.233 | 0.422 | Transketolase, N-terminal |
| IPR020826 | 8.104E-05 | 1.233 | 0.422 | Transketolase binding site |
| IPR001576 | 1.011E-04 | 1.358 | 0.346 | Phosphoglycerate kinase |
| IPR015824 | 1.011E-04 | 1.358 | 0.346 | Phosphoglycerate kinase, N-terminal |
| IPR015901 | 1.011E-04 | 1.358 | 0.346 | Phosphoglycerate kinase, C-terminal |
| IPR015911 | 1.011E-04 | 1.358 | 0.346 | Phosphoglycerate kinase, conserved site |
| IPR010253 | 2.026E-04 | 1.588 | 0.243 | Geranylgeranyl reductase, plant/prokaryotic |
| IPR011774 | 2.026E-04 | 1.588 | 0.243 | Geranylgeranyl reductase, plant/cyanobacteria |
| IPR011777 | 2.026E-04 | 1.588 | 0.243 | Geranylgeranyl reductase family |
| IPR011054 | 2.635E-04 | 1.005 | 0.538 | Rudiment single hybrid motif |
| IPR007066 | 3.756E-04 | 1.071 | 0.461 | RNA polymerase Rpb1, domain 3 |
| IPR007081 | 3.756E-04 | 1.071 | 0.461 | RNA polymerase Rpb1, domain 5 |
| IPR007083 | 3.756E-04 | 1.071 | 0.461 | RNA polymerase Rpb1, domain 4 |
| IPR003960 | 4.785E-04 | 0.633 | 1.165 | ATPase, AAA-type, conserved site |

| | | | | |
|---|---|---|---|---|
| IPR011262 | 4.821E-04 | 1.417 | 0.269 | DNA-directed RNA polymerase, insert domain |
| IPR011263 | 4.821E-04 | 1.417 | 0.269 | DNA-directed RNA polymerase, RpoA/D/Rpb3-type |
| IPR009025 | 4.824E-04 | 1.385 | 0.282 | DNA-directed RNA polymerase, RBP11-like dimerisation domain |
| IPR010004 | 4.894E-04 | 2.154 | 0.128 | Uncharacterised protein family Ycf66 |
| IPR013185 | 4.897E-04 | 1.781 | 0.179 | Translation elongation factor, KOW-like |
| IPR004540 | 4.998E-04 | 1.309 | 0.307 | Translation elongation factor EFG/EF2 |
| IPR020781 | 5.648E-04 | 1.832 | 0.166 | ATPase, OSCP/delta subunit, conserved site |
| IPR000470 | 6.414E-04 | 1.562 | 0.218 | CbxX/CfqX, monofunctional |
| IPR006083 | 6.435E-04 | 1.465 | 0.243 | Phosphoribulokinase/uridine kinase |
| IPR019820 | 6.594E-04 | 1.893 | 0.154 | Sec-independent periplasmic protein translocase, conserved site |
| IPR004176 | 6.633E-04 | 1.151 | 0.371 | Clp, N-terminal |
| IPR018368 | 6.633E-04 | 1.151 | 0.371 | ClpA/B, conserved site 1 |
| IPR028299 | 6.633E-04 | 1.151 | 0.371 | ClpA/B, conserved site 2 |
| IPR019489 | 6.633E-04 | 1.069 | 0.422 | Clp ATPase, C-terminal |
| IPR007072 | 7.824E-04 | 1.584 | 0.205 | Rhamnosyl O-methyltransferase/Cephalosporin hydroxylase |
| IPR011260 | 7.824E-04 | 1.584 | 0.205 | RNA polymerase, alpha subunit, C-terminal |
| IPR011773 | 7.824E-04 | 1.584 | 0.205 | DNA-directed RNA polymerase, alpha subunit |
| IPR000146 | 9.772E-04 | 1.373 | 0.256 | Fructose-1,6-bisphosphatase class 1/Sedoheputulose-1,7-bisphosphatase |
| IPR020548 | 9.772E-04 | 1.373 | 0.256 | Fructose-1,6-bisphosphatase, active site |
| IPR028343 | 9.772E-04 | 1.373 | 0.256 | Fructose-1,6-bisphosphatase |
| IPR002033 | 1.004E-03 | 1.832 | 0.154 | Sec-independent periplasmic protein translocase TatC |
| IPR013785 | 1.041E-03 | 0.468 | 1.818 | Aldolase-type TIM barrel |
| IPR000484 | 1.258E-03 | 1.116 | 0.358 | Photosynthetic reaction centre, L/M |
| IPR007265 | 1.391E-03 | 2.610 | 0.077 | Conserved oligomeric Golgi complex, subunit 3 |
| IPR000568 | 1.405E-03 | 1.417 | 0.230 | ATPase, F0 complex, subunit A |
| IPR023011 | 1.405E-03 | 1.417 | 0.230 | ATPase, F0 complex, subunit A, active site |
| IPR001280 | 1.461E-03 | 1.172 | 0.320 | Photosystem I PsaA/PsaB |
| IPR020586 | 1.461E-03 | 1.172 | 0.320 | Photosystem I PsaA/PsaB, conserved site |
| IPR001270 | 1.659E-03 | 1.012 | 0.410 | ClpA/B family |
| IPR013005 | 1.690E-03 | 1.675 | 0.166 | 50S ribosomal protein uL4 |
| IPR004541 | 1.765E-03 | 1.387 | 0.230 | Translation elongation factor EFTu/EF1A, bacterial/organelle |
| IPR016040 | 1.781E-03 | 0.289 | 4.148 | NAD(P)-binding domain |
| IPR000722 | 1.951E-03 | 1.169 | 0.307 | RNA polymerase, alpha subunit |
| IPR006592 | 1.951E-03 | 1.169 | 0.307 | RNA polymerase, N-terminal |
| IPR007080 | 1.951E-03 | 1.169 | 0.307 | RNA polymerase Rpb1, domain 1 |
| IPR002325 | 2.393E-03 | 1.626 | 0.166 | Cytochrome f |
| IPR024058 | 2.393E-03 | 1.626 | 0.166 | Cytochrome f transmembrane anchor |
| IPR003672 | 3.068E-03 | 1.247 | 0.256 | CobN/magnesium chelatase |
| IPR011771 | 3.068E-03 | 1.247 | 0.256 | Magnesium-chelatase, subunit H |
| IPR022571 | 3.068E-03 | 1.247 | 0.256 | Magnesium chelatase, subunit H, N-terminal |
| IPR004853 | 3.131E-03 | 1.101 | 0.320 | Sugar phosphate transporter domain |
| IPR001263 | 3.346E-03 | 1.454 | 0.192 | Phosphoinositide 3-kinase, accessory (PIK) domain |
| IPR023222 | 3.346E-03 | 1.454 | 0.192 | PsbQ-like domain |
| IPR024094 | 3.365E-03 | 1.578 | 0.166 | Cytochrome f large domain |
| IPR010251 | 3.842E-03 | 2.417 | 0.077 | Magnesium-protoporphyrin IX methyltransferase |
| IPR010940 | 3.842E-03 | 2.417 | 0.077 | Magnesium-protoporphyrin IX methyltransferase, C-terminal |
| IPR002358 | 4.366E-03 | 1.417 | 0.192 | Ribosomal protein L6, conserved site |
| IPR019906 | 4.366E-03 | 1.417 | 0.192 | Ribosomal protein L6, bacterial-type |
| IPR000711 | 4.386E-03 | 1.533 | 0.166 | ATPase, OSCP/delta subunit |
| IPR005869 | 4.386E-03 | 1.533 | 0.166 | Photosystem II CP43 reaction centre protein |
| IPR026015 | 4.386E-03 | 1.533 | 0.166 | F1F0 ATP synthase OSCP/delta subunit, N-terminal domain |
| IPR005493 | 4.407E-03 | 2.180 | 0.090 | Ribonuclease E inhibitor RraA/Dimethylmenaquinone methyltransferase |
| IPR022147 | 4.428E-03 | 1.610 | 0.154 | Glutamine synthetase type III N-terminal |
| IPR000641 | 4.466E-03 | 1.274 | 0.230 | CbxX/CfqX |
| IPR005706 | 4.474E-03 | 1.142 | 0.282 | Ribosomal protein S2, bacteria/mitochondria/plastid |
| IPR018936 | 7.415E-03 | 1.116 | 0.269 | Phosphatidylinositol 3/4-kinase, conserved site |
| IPR000523 | 7.765E-03 | 1.695 | 0.128 | Magnesium chelatase ChlI domain |
| IPR010136 | 7.905E-03 | 2.832 | 0.051 | N-acetyl-gamma-glutamyl-phosphate reductase, type 2 |
| IPR000652 | 9.706E-03 | 1.008 | 0.307 | Triosephosphate isomerase |
| IPR020861 | 9.706E-03 | 1.008 | 0.307 | Triosephosphate isomerase, active site |
| IPR009014 | 9.807E-03 | 0.794 | 0.474 | Transketolase, C-terminal/Pyruvate-ferredoxin oxidoreductase, domain II |

**Supplementary Table S8.** Enriched InterPro domains in diatom-assigned transcripts expressed in the Wilkins sea ice (WKI) sample, compared with the whole Antarctic Peninsula phytoplankton community metatranscriptome. Enrichment fold values are expressed in $\log_2$ scale and the subset ratio indicates the fraction of transcripts annotated with the enriched functional annotation label within the subset. Maximum enrichment *q*-value threshold is 0.01.

| Identifier | *q*-value | Enrichment fold | Subset ratio | Description |
|---|---|---|---|---|
| IPR022796 | 5.235E-37 | 0.990 | 3.778 | Chlorophyll A-B binding protein |
| IPR023329 | 1.452E-36 | 0.975 | 3.810 | Chlorophyll a/b binding protein domain |
| IPR003593 | 2.960E-08 | 0.425 | 4.504 | AAA+ ATPase domain |
| IPR004737 | 1.271E-07 | 2.389 | 0.160 | Nitrate transporter |
| IPR003959 | 3.236E-07 | 0.633 | 1.868 | ATPase, AAA-type, core |
| IPR005936 | 5.532E-07 | 1.260 | 0.491 | Peptidase, FtsH |
| IPR005117 | 7.081E-07 | 1.649 | 0.299 | Nitrite/Sulfite reductase ferredoxin-like domain |
| IPR006066 | 7.081E-07 | 1.649 | 0.299 | Nitrite/sulphite reductase iron-sulphur/sirohaem-binding site |
| IPR006067 | 7.081E-07 | 1.649 | 0.299 | Nitrite/sulphite reductase 4Fe-4S domain |
| IPR000642 | 7.815E-07 | 1.223 | 0.491 | Peptidase M41 |
| IPR000185 | 8.533E-07 | 1.448 | 0.363 | Protein translocase subunit SecA |
| IPR011115 | 8.533E-07 | 1.448 | 0.363 | SecA DEAD-like, N-terminal |
| IPR011116 | 8.533E-07 | 1.448 | 0.363 | SecA Wing/Scaffold |
| IPR011130 | 8.533E-07 | 1.448 | 0.363 | SecA preprotein, cross-linking domain |
| IPR014018 | 8.533E-07 | 1.448 | 0.363 | SecA motor DEAD |
| IPR020937 | 8.533E-07 | 1.448 | 0.363 | SecA conserved site |
| IPR017871 | 1.015E-06 | 0.603 | 1.847 | ABC transporter, conserved site |
| IPR003439 | 1.168E-06 | 0.576 | 1.932 | ABC transporter-like |
| IPR021884 | 4.504E-06 | 1.136 | 0.502 | Protein of unknown function DUF3494 |
| IPR003960 | 5.965E-06 | 0.695 | 1.217 | ATPase, AAA-type, conserved site |
| IPR006073 | 6.141E-06 | 0.888 | 0.768 | GTP binding domain |
| IPR017926 | 7.433E-05 | 0.918 | 0.598 | Glutamine amidotransferase |
| IPR027417 | 7.474E-05 | 0.181 | 12.008 | P-loop containing nucleoside triphosphate hydrolase |
| IPR007419 | 1.876E-04 | 2.122 | 0.117 | BFD-like [2Fe-2S]-binding domain |
| IPR012744 | 1.876E-04 | 2.122 | 0.117 | Nitrite reductase [NAD(P)H] large subunit, NirB |
| IPR003672 | 3.972E-04 | 1.307 | 0.267 | CobN/magnesium chelatase |
| IPR011771 | 3.972E-04 | 1.307 | 0.267 | Magnesium-chelatase, subunit H |
| IPR022571 | 3.972E-04 | 1.307 | 0.267 | Magnesium chelatase, subunit H, N-terminal |
| IPR029062 | 5.998E-04 | 0.790 | 0.640 | Class I glutamine amidotransferase-like |
| IPR002792 | 6.225E-04 | 1.570 | 0.181 | TRAM domain |
| IPR000183 | 6.912E-04 | 1.669 | 0.160 | Ornithine/DAP/Arg decarboxylase |
| IPR010253 | 8.916E-04 | 1.400 | 0.213 | Geranylgeranyl reductase, plant/prokaryotic |
| IPR011774 | 8.916E-04 | 1.400 | 0.213 | Geranylgeranyl reductase, plant/cyanobacteria |
| IPR011777 | 8.916E-04 | 1.400 | 0.213 | Geranylgeranyl reductase family |
| IPR005706 | 9.673E-04 | 1.175 | 0.288 | Ribosomal protein S2, bacteria/mitochondria/plastid |
| IPR011545 | 9.749E-04 | 0.435 | 1.825 | DEAD/DEAH box helicase domain |
| IPR012748 | 9.816E-04 | 1.833 | 0.128 | Nitrite reductase [NAD(P)H] small subunit, NirD |
| IPR014001 | 9.965E-04 | 0.390 | 2.199 | Helicase superfamily 1/2, ATP-binding domain |
| IPR001054 | 1.020E-03 | 1.062 | 0.342 | Adenylyl cyclase class-3/4/guanylyl cyclase |
| IPR002073 | 1.025E-03 | 1.102 | 0.320 | 3'5'-cyclic nucleotide phosphodiesterase, catalytic domain |
| IPR003029 | 1.026E-03 | 0.830 | 0.534 | S1 domain |
| IPR012756 | 1.182E-03 | 1.318 | 0.224 | DNA-directed RNA polymerase, subunit beta" |
| IPR020547 | 1.192E-03 | 1.985 | 0.107 | ATP synthase delta/epsilon subunit, C-terminal domain |
| IPR029787 | 1.219E-03 | 1.046 | 0.342 | Nucleotide cyclase |
| IPR001650 | 1.317E-03 | 0.384 | 2.156 | Helicase, C-terminal |
| IPR022967 | 1.327E-03 | 0.813 | 0.534 | RNA-binding domain, S1 |
| IPR022643 | 1.360E-03 | 1.570 | 0.160 | Orn/DAP/Arg decarboxylase 2, C-terminal |
| IPR022644 | 1.360E-03 | 1.570 | 0.160 | Orn/DAP/Arg decarboxylase 2, N-terminal |
| IPR014102 | 1.478E-03 | 1.685 | 0.139 | Phytoene desaturase |

| | | | | |
|---|---|---|---|---|
| IPR007148 | 1.525E-03 | 1.448 | 0.181 | Small-subunit processome, Utp12 |
| IPR020846 | 1.690E-03 | 0.542 | 1.089 | Major facilitator superfamily domain |
| IPR007034 | 1.833E-03 | 1.382 | 0.192 | Ribosome biogenesis protein BMS1/TSR1, C-terminal |
| IPR012948 | 1.833E-03 | 1.382 | 0.192 | AARP2CN |
| IPR009006 | 1.916E-03 | 1.522 | 0.160 | Alanine racemase/group IV decarboxylase, C-terminal |
| IPR023179 | 2.104E-03 | 1.222 | 0.235 | GTP-binding protein, orthogonal bundle domain |
| IPR000232 | 2.345E-03 | 1.155 | 0.256 | Heat shock factor (HSF)-type, DNA-binding |
| IPR004693 | 2.443E-03 | 1.440 | 0.171 | Silicon transporter |
| IPR012137 | 4.120E-03 | 1.804 | 0.107 | Nitrate reductase NADH dependent |
| IPR002937 | 4.606E-03 | 1.002 | 0.299 | Amine oxidase |
| IPR018130 | 4.706E-03 | 0.860 | 0.395 | Ribosomal protein S2, conserved site |
| IPR023591 | 4.706E-03 | 0.860 | 0.395 | Ribosomal protein S2, flavodoxin-like domain |
| IPR001138 | 4.777E-03 | 2.347 | 0.064 | Zn(2)-C6 fungal-type DNA-binding domain |
| IPR000511 | 4.856E-03 | 2.570 | 0.053 | Cytochrome c/c1 haem-lyase |
| IPR003728 | 4.856E-03 | 2.570 | 0.053 | Ribosome maturation factor RimP |
| IPR004154 | 5.558E-03 | 0.857 | 0.384 | Anticodon-binding |
| IPR009025 | 5.565E-03 | 1.122 | 0.235 | DNA-directed RNA polymerase, RBP11-like dimerisation domain |
| IPR011262 | 5.576E-03 | 1.155 | 0.224 | DNA-directed RNA polymerase, insert domain |
| IPR011263 | 5.576E-03 | 1.155 | 0.224 | DNA-directed RNA polymerase, RpoA/D/Rpb3-type |
| IPR002146 | 5.661E-03 | 1.300 | 0.181 | ATPase, F0 complex, subunit B/B', bacterial/chloroplast |
| IPR002033 | 5.717E-03 | 1.570 | 0.128 | Sec-independent periplasmic protein translocase TatC |
| IPR005717 | 6.136E-03 | 1.833 | 0.096 | Ribosomal protein S7, bacterial/organellar-type |
| IPR013816 | 6.338E-03 | 0.560 | 0.822 | ATP-grasp fold, subdomain 2 |
| IPR030378 | 6.518E-03 | 1.164 | 0.213 | Circularly permuted (CP)-type guanine nucleotide-binding (G) domain |
| IPR002938 | 6.902E-03 | 0.863 | 0.363 | FAD-binding domain |
| IPR011701 | 6.976E-03 | 0.685 | 0.555 | Major facilitator superfamily |
| IPR023753 | 7.049E-03 | 0.384 | 1.633 | FAD/NAD(P)-binding domain |
| IPR008333 | 8.271E-03 | 1.137 | 0.213 | Oxidoreductase, FAD-binding domain |
| IPR011060 | 8.360E-03 | 0.821 | 0.384 | Ribulose-phosphate binding barrel |
| IPR011761 | 8.566E-03 | 0.580 | 0.726 | ATP-grasp fold |
| IPR003607 | 9.285E-03 | 0.950 | 0.288 | HD/PDEase domain |
| IPR010073 | 9.304E-03 | 1.570 | 0.117 | Phosphoribosylformylglycinamidine synthase |
| IPR022653 | 9.603E-03 | 1.869 | 0.085 | Orn/DAP/Arg decarboxylase 2, pyridoxal-phosphate binding site |
| IPR022657 | 9.603E-03 | 1.869 | 0.085 | Orn/DAP/Arg decarboxylase 2, conserved site |
| IPR013005 | 9.694E-03 | 1.412 | 0.139 | 50S ribosomal protein uL4 |
| IPR010004 | 9.732E-03 | 1.740 | 0.096 | Uncharacterised protein family Ycf66 |
| IPR002986 | 9.738E-03 | 1.644 | 0.107 | Diaminopimelate decarboxylase, LysA |
| IPR007109 | 9.809E-03 | 1.110 | 0.213 | Brix domain |
| IPR001865 | 9.829E-03 | 0.816 | 0.374 | Ribosomal protein S2 |

**Supplementary Table S9.** 'Minimal' taxonomic classification evaluation dataset composition and sampling proportions. Sequence counts indicate the number of available RNA sequences from each sampled clade in RefSeq (release 84).

| Clade (NCBI taxon identifier) | Rank | #sequences | Fraction |
|---|---|---|---|
| Chlorophyta (3041) | Phylum | 115,801 | 0.20 |
| Streptophyta (35493) | Phylum | 4,190,204 | 0.20 |
| Bacteria (2) | Superkingdom | 21,190 | 0.15 |
| Chordata (7711) | Phylum | 9,494,012 | 0.05 |
| Basidiomycota (5204) | Phylum | 486,296 | 0.05 |
| Arthropoda (6656) | Phylum | 2,524,444 | 0.05 |
| Ascomycota (4890) | Phylum | 1,685,068 | 0.05 |
| Mollusca (6447) | Phylum | 218,941 | 0.05 |
| Apicomplexa (5794) | Phylum | 186,627 | 0.05 |
| Nematoda (6231) | Phylum | 166,442 | 0.05 |
| Cnidaria (6073) | Phylum | 154,324 | 0.05 |
| Archaea (2157) | Superkingdom | 1,162 | 0.05 |
